# Degrading intestinal DAF-2 nearly doubles *Caenorhabditis elegans* lifespan without affecting development or reproduction

**DOI:** 10.1101/2021.07.31.454567

**Authors:** Yan-Ping Zhang, Wen-Hong Zhang, Pan Zhang, Qi Li, Yue Sun, Jia-Wen Wang, Shao-Bing O. Zhang, Tao Cai, Cheng Zhan, Meng-Qiu Dong

## Abstract

Twenty-eight years following the breakthrough discovery that a single-gene mutation of *daf-2* can double the lifespan of *Caenorhabditis elegans*, it remains unclear where this gene, which encodes an insulin/IGF-1 receptor, is expressed and where it acts to regulate aging. Here, by inserting DNA sequences of fluorescent tags into the genomic locus of *daf-2* and that of its downstream transcription factor *daf-16*, we determined that both genes are expressed in most or all tissues from embryos through adulthood, in line with their diverse functions. Using tissue-specific auxin-induced protein degradation, we determined that both DAF-2 and DAF-16 act in the intestine to regulate organismal aging. Strikingly, loss of DAF-2 in the intestine nearly doubled *C. elegans* lifespan but did not produce the adverse developmental or reproductive phenotypes associated with genetic *daf-2* mutants. These findings unify the mechanism of lifespan regulation by genes and that by dietary restriction, and begin to focus anti-aging research on nutrient supply.

**Highlights:**

1. *daf-2* and *daf-16* are expressed in most or all cells of *C. elegans* using genome editing.
2. DAF-2 and DAF-16 both regulate lifespan from the intestine as determined using auxin-induced protein degradation.
3. Reduced insulin signaling in the intestine nearly doubles *C. elegans* lifespan without adverse effects on development or reproduction.
4. Lifespan regulation by genes and dietary restriction are unified by intestinal supply of nutrients and metabolism.

## Introduction

One of the breakthrough discoveries in biology from the last 30 years is the finding that ancient genetic pathways control animal lifespan (Kenyon, 2001; Kenyon, 2010; Narasimhan et al., 2009). The first identified and extensively validated one is the insulin/insulin-like growth factor 1 (IGF-1) signaling (IIS) pathway (Guarente and Kenyon, 2000; Kenyon, 2011; Kenyon et al., 1993). Reduction of the IIS extends the lifespan in *Caenorhabditis elegans* (Friedman and Johnson, 1988; Kenyon et al., 1993; Morris et al., 1996), *Drosophila* (Clancy et al., 2001; Tatar et al., 2001), and mice (Bluher et al., 2003; Holzenberger et al., 2003). Moreover, single nucleotide polymorphisms (SNPs) of IIS component genes are tightly linked to human longevity (Pawlikowska et al., 2009; Suh et al., 2008; Willcox et al., 2008). Reducing IIS also alleviates pathologies of age-associated diseases in animal models, including those for Alzheimer’s and Parkinson’s (Cohen et al., 2009; El-Ami et al., 2014). These studies indicate that the IIS pathway is a promising target for developing anti-aging therapeutics. However, applying this knowledge in practice has not been possible because of the many essential functions of IIS, including glucose metabolism, lipid metabolism, growth, and reproduction (Barbieri et al., 2003; Piper et al., 2008; Saltiel and Kahn, 2001; Zhang and Liu, 2014).

In humans and other mammals, insulin and IGF peptides are synthesized in and secreted primarily from the pancreatic β-cells and the liver, respectively. They are carried by the bloodstream to target tissues where they bind to and activate their cell-surface receptors. In humans, both the insulin receptor (IR) and the IGF-1 receptor (IGF-1R) are expressed in nearly all tissues (https://portal.brain-map.org/). In mammals, studies of IIS mostly focus on β-cells, liver, muscle, and adipose tissue, where IIS is crucial in maintaining homeostasis of glucose metabolism and energy metabolism (Belfiore et al., 2017; Zhang and Liu, 2014). Given these essential functions, mutations of IRs are linked to several inheritable genetic diseases including type A insulin resistance syndrome, Donohue syndrome, and Rabson-Mendenhall syndrome (Plamper et al., 2018). In healthy people, sensitivity to insulin declines with age, accompanied by an increasing probability of developing serious chronic conditions such as type 2 diabetes and obesity (Boucher et al., 2014; Czech, 2017; Titchenell et al., 2017). As such, it was surprising when reduction of IIS was discovered to extend lifespan of *C. elegans* (Kenyon et al., 1993). Interestingly, some forms of general IIS reduction are accompanied by a longevity phenotype in humans, mice, and dogs, although they also exhibit an undesirable growth retardation phenotype (Kenyon, 2010).

The *daf-2* gene encodes the sole *C. elegans* homolog of IR/IGF-1R (Kimura et al., 1997). Other core components of IIS include AGE-1/PI3-K, PDK-1, AKT-1/2, and DAF-16/FoxO (Kimura et al., 1997; Lin et al., 1997; Ogg et al., 1997; Paradis and Ruvkun, 1998; Pierce et al., 2001). The IIS kinase cascade, from DAF-2 to AKT-1/2, maintains a relatively short wild-type (WT) lifespan by inhibiting the transcription factor (TF) DAF-16. Loss-of-function (*lf*) mutations of the upstream kinases all produce a remarkable longevity phenotype in a *daf-16* dependent manner. For example, the canonical *daf-2(e1370)* allele doubles *C. elegans* lifespan, and the lifespan extension is completely abolished by deletion of *daf-16* (Kenyon et al., 1993). Reduction of *C. elegans* IIS from young adulthood produces a stronger longevity phenotype than reducing it in later periods (Dillin et al., 2002). In addition to a long lifespan, similar to mammals, *C. elegans* IIS mutants are highly pleiotropic, exhibiting varying degrees of developmental defects, reduced brood size, and increased fat storage (Gems et al., 1998; Kimura et al., 1997).

Various methods, from genetic mosaic analysis (Apfeld and Kenyon, 1998) to tissue-specific RNAi (Uno et al., 2021) or transgene rescue (Libina et al., 2003), have been tried to identify the different functions of IIS in *C. elegans*. Special emphasis has been placed on determining the tissue(s) from which IIS regulates lifespan, partly because this question has significant implications in itself, and partly because *C. elegans* is an excellent model for aging research. However, despite these efforts, no consensus has been reached: experimental data have suggested neurons (Wolkow et al., 2000), intestine (Libina et al., 2003), or multiple cell lineages (Apfeld and Kenyon, 1998) as the sites of lifespan regulation by IIS.

The main source of the confusion is that the exact expression pattern of *daf-2* is unknown. The *daf-2* gene is 50 kb long, with large introns, multiple transcription start sites and alternative splicing sites; there are also multiple, long cDNAs. These characteristics make it difficult to determine the expression pattern of *daf-2* with traditional transgene approaches. Immunostaining and *in situ* hybridization did not produce consistent results, either, with one showing DAF-2 in XXX cells and neurons (Kimura et al., 2011) and the other showing DAF-2 in the germline (Honnen et al., 2012). In contrast, independent studies using fluorescent transgene reporters all agree that *daf-16* is ubiquitously expressed in somatic tissues (Henderson and Johnson, 2001; Kwon et al., 2010; Lee et al., 2001; Lin et al., 2001).

In this study, using CRISPR/Cas9 genome editing technology and tissue-specific targeted protein degradation system, we determined the endogenous expression patterns of *daf-2* and *daf-16* and their sites of action in lifespan regulation. We found that DAF-2 and DAF-16 are both expressed ubiquitously in the somatic and reproductive tissues, and that both regulate aging of the entire body from the intestine without interfering with development and reproduction. Further, degrading intestinal DAF-2 nearly doubles *C. elegans* lifespan. These findings suggest that genetic regulation of aging by IIS operates by affecting nutrient supply from the intestine to the entire body, thus inducing metabolic changes at the organismal level. The evolutionarily conserved lifespan regulating mechanisms including IIS and Mtor signaling all sense nutrients. Hence, we postulate that genetic regulation of aging as well as dietary regulation of aging of various forms including caloric restriction, dietary restriction, and intermittent feeding, must converge on the downstream metabolic pathways. Our data suggest that down-regulation of protein synthesis is a molecular signature of longevity. These findings provide insights that unify the superficially diverse anti-aging mechanisms and point a way to achieving longevity without adverse effects in development and reproduction.

## Results

### 1. Insulin/IGF-1 receptor, a master regulator of aging, is widely expressed in *C. elegans* from embryos to adults

To ensure accurate detection of the endogenous expression pattern of *daf-2*, we designed two detection strategies, one focusing on the protein, the other on the mRNA (Figure 1A).

**Figure 1.**
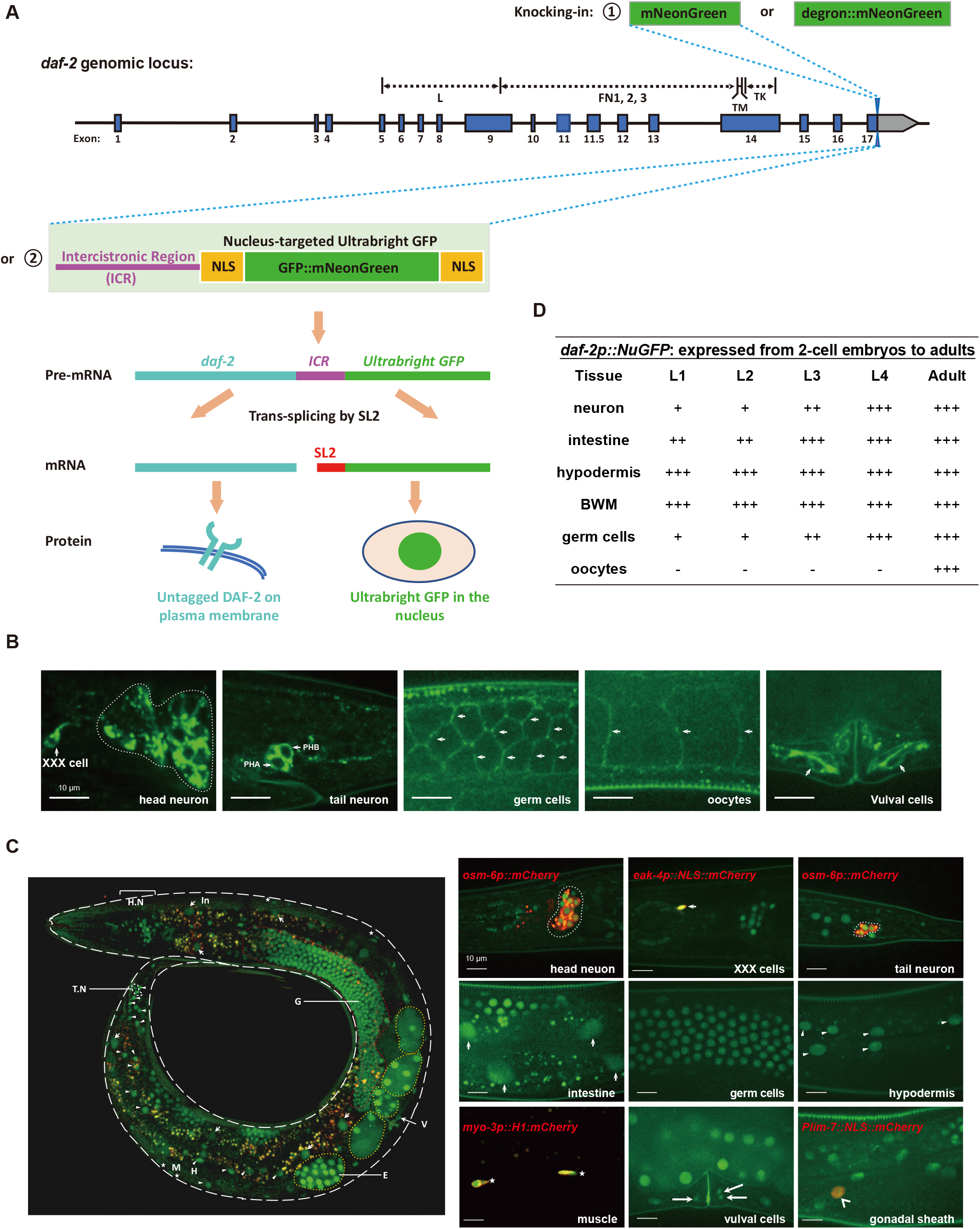
Endogenous expression of the *daf-2* gene detected in most or all *C. elegans* cells from embryos to adults. (A) Schematic of two strategies to characterize the endogenous expression pattern of *daf-2*. The coding sequence of the mNeonGreen, mNeonGreen::degron (top panel), or NuGFP cassette (bottom panel) is knocked into the *daf-2* genomic locus before the stop codon by CRISPR/Cas9 genome editing, which allows detection of *daf-2* expression at the protein level or at the mRNA level, respectively. Blue boxes, coding regions; line, non-coding regions; grey boxes, 3’ untranslated regions. (B) Expression pattern of *daf-2* at the protein level indicated by the DAF-2::mNeonGreen at day 1 of adulthood. (C) Expression pattern of *daf-2* at the mRNA level indicated by the NuGFP reporter at day 1 of adulthood. Left panel, overview of the expression of *daf-2* mRNA throughout the whole body. H.N, head neuron; G, germ line, indicated by the circled red dotted line; In, intestine, indicated by short arrows; V, vulval cells, indicated by long arrow; E, embryo, indicated by the circled grey dashed line; H, hypodermal cells, indicated by triangles; T.N, tail neuron, indicated by the circled white dotted line; M, body wall muscle, indicated by asterisks. Right panel, local expression patterns of *daf-2* mRNA. *osm-6p::mCherry*, ciliated sensory neuron marker; *eak-4p::NLS::mCherry*, XXX cells-specific marker; *myo-3p::H1::mCherry*, body wall muscle-specific marker; *lim-7p::NLS::mCherry*, gonadal sheath-specific marker. (D) Summary of the spatiotemporal expression pattern of NuGFP reporter. +, expression intensity of NuGFP; -, expression of NuGFP is not detectable. See also Figures S1 and S2.

To visualize the DAF-2 protein, we knocked in the CDS of mNeonGreen, a green fluorescent protein four times as bright as the jellyfish GFP (Shaner et al., 2013), immediately after the last amino acid codon of the *daf-2* gene on chromosome III (Figure 1A). Phenotypic assays showed that this mNeonGreen tag did not perturb the function of the DAF-2 protein to which it was attached (Figure S1). DAF-2::mNeonGreen was detected in neurons, XXX cells, vulval cells, germ cells, and oocytes (Figure 2B). In the last three types of cells, DAF-2::mNeonGreen had clear plasma membrane (PM) localization as expected for a cell surface receptor. In the neurons and XXX cells (Figure 1B), a pair of specialized hypodermal cells of neural endocrine function, strong DAF-2::mNeonGreen signals were seen in the cell bodies and along processes. In addition, the fluorescence signal appeared to originate from both the PM and the cytoplasm, presumably from intracellular membranes along the synthesis and secretion pathways of proteins targeted to the PM. Presence of DAF-2 in vulval cells was not suggested before, but Nadkimon et al. (2012) showed that *daf-2(lf)* suppressed the multivulva phenotype induced by hyperactivation of RAS/MAPK signaling (Nakdimon et al., 2012). Therefore, the above results indicate a cell-autonomous regulatory function of IIS in vulval development. The DAF-2::mNeonGreen-expressing neurons included all of the ciliated sensory neurons marked by the *osm-6* promoter-driven mCherry protein (Figure S2A).

**Figure 2.**
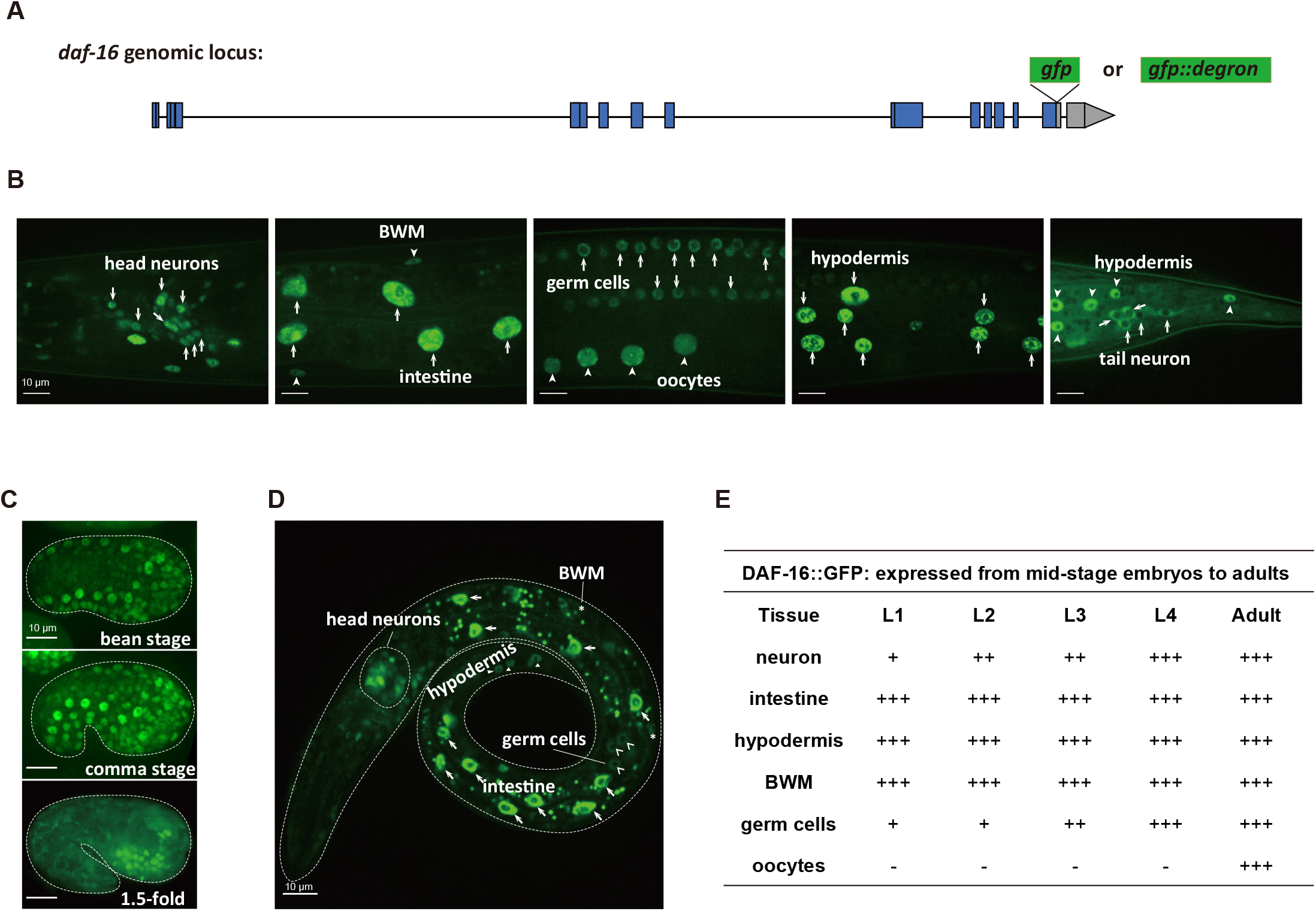
Ubiquitous presence of DAF-16 detected in the *C. elegans* soma and germline. (A) Schematic of knocking in the coding sequence of GFP or GFP::degron into the *daf-16* genomic locus before the stop codon using CRISPR/Cas9 genome editing. Blue boxes, coding regions; line, non-coding regions; grey boxes, 3’ untranslated regions. (B-D) Expression patterns of DAF-16::GFP at day 1 of adulthood (B), embryonic stage (C), and L1 larval stage (D). (E) Summary of the spatiotemporal expression patterns of DAF-16::GFP. +, expression intensity of DAF-16::GFP; -, expression of DAF-16::GFP is not detectable. See also Figures S1 and S3.

Because DAF-2 is localized on the PM, it may be difficult to detect using the mNeonGreen tag in cells with large surface area where the signal may be too thin. Therefore, we took an alternative approach to visualize the cells expressing *daf-2* mRNA. Downstream of the *daf-2* CDS, we knocked in the sequence of an intercistronic region (ICR) from the *C. elegans* SL2-type operon, followed by a nuclear localization sequence (NLS), the CDS of GFP, the CDS of mNeonGreen, and another NLS, which we referred to as the Nuclear Ultrabright GFP::mNeonGreen Fluorescent protein (NuGFP) cassette (Figure 1A). In this approach, expression of NuGFP is tied to that of *daf-2* from the endogenous locus, but after trans-splicing, the NuGFP protein is synthesized independently of DAF-2. This high-sensitivity *daf-2* expression reporter was readily detectable in most *C. elegans* cells, including the ones that had been missed by the DAF-2::mNeonGreen fusion protein marker, that is, the intestine, hypodermis, gonadal sheath, and BWM (Figure 1C). With NuGFP, expression of *daf-2* was observed starting in 2-cell embryos, and the expression continued throughout development and adulthood (Figure 1D and Figure S2B-S2G).

### 2. The FoxO transcription factor is widely expressed in *C. elegans* from embryos to adults

To visualize the pattern of endogenous expression of *daf-16*, we knocked in the coding sequence (CDS) of the green fluorescent protein (GFP) right before the stop codon of the *daf-16* CDS on chromosome III using the CRISPR/Cas9 genome editing technique (Figure 2A). Other than illuminating the endogenously expressed DAF-16 proteins, the GFP tag did not interfere with DAF-16 function (Figure S1).

In unstressed animals, such as those kept at standard culture conditions (15-20 °C, well-fed), DAF-16::GFP was dispersed throughout the cell. To better recognize cells expressing DAF-16::GFP, we induced nuclear translocation of DAF-16::GFP by placing the worms on a glass slide atop an agarose cushion for about 5 minutes before epifluorescence imaging. We found that DAF-16::GFP was expressed ubiquitously in most or all somatic tissues, such as neurons, intestine, body wall muscles (BWM), and hypodermis, and also in the germ cells and oocytes (Figure 2B). Germline expression of DAF-16::GFP was not detected by earlier transgene reporters (Henderson and Johnson, 2001; Kwon et al., 2010; Lee et al., 2001; Lin et al., 2001). Temporally, ubiquitous expression of DAF-16::GFP from the endogenous locus started during embryonic development at the bean stage and persisted through larval development and adulthood (Figure 2C-2E and Figure S3).

### 3. Insulin/IGF-1 receptor controls lifespan from the intestine

To settle the controversy regarding the site of action of insulin signaling in lifespan regulation, we adopted the auxin-induced protein degradation (AID) system (Zhang et al., 2015) (Figure S4A). This system allowed us to achieve tissue-specific DAF-2 or DAF-16 degradation in the WT or the long-lived *daf-2(e1370)* mutant background, respectively.

Using CRISPR/Cas9 technology, we generated knocking-in strains respectively expressing DAF-2::degron::mNeonGreen (Figure 1A) or DAF-16::GFP::degron (Figure 2A) from the endogenous *daf-2* or *daf-16* locus. The double tag of a degron sequence and a fluorescent protein sequence enables facile detection of the expression of the fusion protein and auxin-induced target protein degradation. Next, to the existing single-copy insertion (SCI) strains expressing TIR-1 in all cells (*ieSi57*), intestinal cells (*ieSi61*), or germ cells (*ieSi38*), we added neuron-, hypodermis-, BWM-, gonadal-sheath-, and XXX-cell-specific TIR-1 expressing lines by replacing the promoter sequence of *eft-3* or *sun-1* with that of *rgef-1, dpy-7, myo-3, lim-7*, or *eak-4* (Figure S4B). The above TIR-1 expressing chromosomes II or IV were each combined with the DAF-2::degron::mNeonGreen or DAF-16::GFP::degron chromosome through genetic crossing. Auxin-induced tissue-specific degradation of fluorescently labeled DAF-2 or DAF-16 was verified (Figure S4C-S4J and Figure S5).

Next, we examined the effect of tissue-specific degradation of DAF-2 on lifespan (Figure 3 and Table S1). We found that degrading neuronal DAF-2 increased WT lifespan by 18.6% (Figure 3A), much less than what would be expected from previous *daf-2(lf)* rescue experiments using the tissue-specific-promoter-driven transgene arrays (Wolkow et al., 2000). Degrading DAF-2 in the germline or the hypodermis respectively increased lifespan by 6.4% and 13.7% (Figure 3B and 3C), whereas degrading DAF-2 in the BWM, gonadal sheath, or XXX cells had no effect on lifespan (Figure 3D-3F). In contrast, degrading intestinal DAF-2 extended the *C. elegans* lifespan by 94.3% (Figure 3G). The above results showed unambiguously that intestinal DAF-2 is essential in lifespan regulation; neuronal, hypodermal, and germline DAF-2 each play a minor role, while DAF-2 has no detectable effect in other tissues.

**Figure 3.**
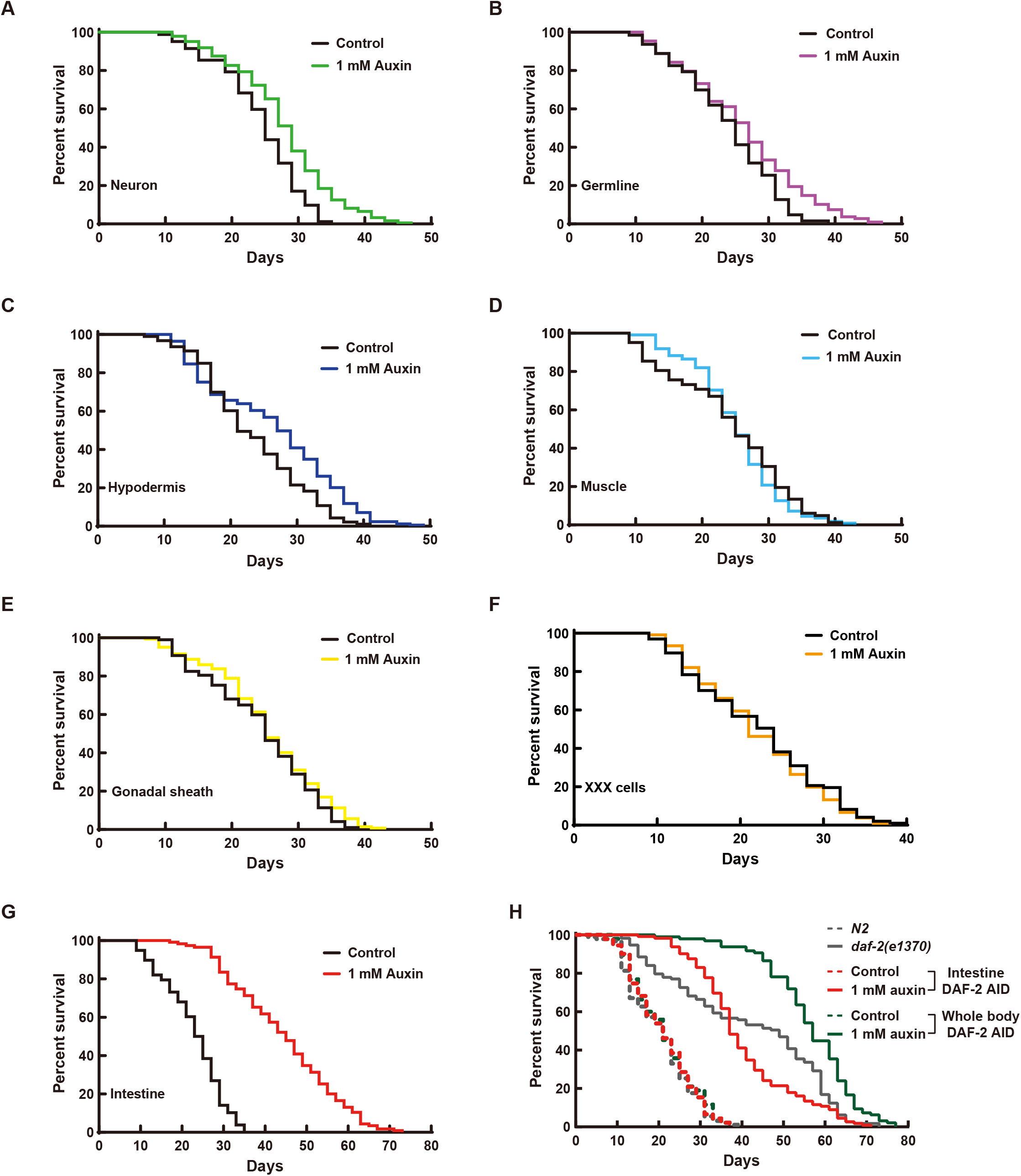
Intestine-specific degradation of DAF-2 extended *C. elegans* lifespan by 94.3%. (A-C) Degrading DAF-2 in the neurons (A), germline (B), and hypodermis (C) increased the lifespan by 18.6% (*p*<0.0001), 6.4% (*p*<0.05), and 13.7% (*p*<0.001), respectively. (D-F) Degrading DAF-2 in the BWM (D), gonadal sheath (E), or XXX cells (F) had no effect on WT lifespan (*p*>0.05). (G) Degrading DAF-2 in the intestine increased WT lifespan by 94.3% (*p*<0.0001). (H) Degrading DAF-2 in the whole body increased WT lifespan by 166.5% (green solid line versus green dashed line, *p*<0.0001), which outlived the canonical hypomophic *daf-2(e1370)* mutant worms and the intestinal DAF-2 AID worms (grey solid line and red solid line, respectively). *p*-values are calculated by log-rank tests. See also Table S1 and Figure S4.

Of note, the worms in which DAF-2 was degraded throughout the body had a lifespan that was 266.5% of the WT (Figure 3H), outliving the canonical hypomophic *daf-2(e1370)* mutant and the intestinal DAF-2 AID worms, whose lifespans were 205.9% and 192.1% of the WT, respectively (Figure 3H). Here, FUdR was used to prevent the strong egg-laying defective (Egl) phenotype of whole-body DAF-2 AID from interfering with the lifespan assay. A strong Egl phenotype causes internal hatching of eggs. Degrading DAF-2 in the intestine or the other tissues (Figure 3A-3G) caused no obvious Egl phenotype.

The data above argue unequivocally that the insulin/IGF-1 receptor regulates aging of the entire body from the intestine, not neurons. This has implications beyond clarifying the purported role of neuronal DAF-2 in lifespan regulation (Wolkow, 2000), which has generated both excitement and uncertainty. Given that the intestine supplies nutrients to the entire body and insulin signaling regulates metabolism (Saltiel and Kahn, 2001; Zhang and Liu, 2014), it makes sense that IIS controls animal aging from an organ or tissue that is the center of nutrient supply. In addition, the above results suggest that lifespan regulation by genes and by dietary restriction likely share common downstream mechanisms. The clear lifespan extension (whole-body DAF-2 AID 166.5% > intestine 94.3% + neuron 18.6% + hypodermis 13.7% + germline 6.4% + others 0%) is a reminder that aging is a systems phenomenon involving cooperation among different tissues.

### 4. Intestinal FoxO mediates lifespan extension by reduced insulin/IGF-1 signaling

The longevity phenotype of *daf-2(e1370)* is fully dependent on *daf-16* (Kenyon et al., 1993), therefore one obvious question is in which tissue DAF-16 mediates this effect. Degrading DAF-16 in the *daf-2(e1370)* background in a tissue-specific manner (Figure 4 and Table S1) in the neurons, germline, or hypodermis shortened the *daf-2(e1370)* lifespan by no more than 15.6% (Figure 4A-4C), while degrading DAF-16 in BWM, gonadal sheath, or XXX cells had no effect (Figure 4D-4F). In comparison, degrading intestinal DAF-16 shortened the *daf-2(e1370)* lifespan by 40.1% (Figure 4G), which means that intestinal DAF-16 mediated 90.3% of the lifespan extension by *daf-2(e1370)*. Degrading DAF-16 throughout the body shortened the *daf-2(e1370)* lifespan by 57.6%, making it slightly shorter than the lifespan of WT animals at 44.4% of the *daf-2(e1370)* lifespan (Figure 4H). The effects of tissue-specific DAF-16 degradation resonated those of tissue-specific DAF-2 degradation on WT lifespan (Figure 3), and clearly indicate that the intestine is the single most important tissue from which IIS regulates lifespan.

**Figure 4.**
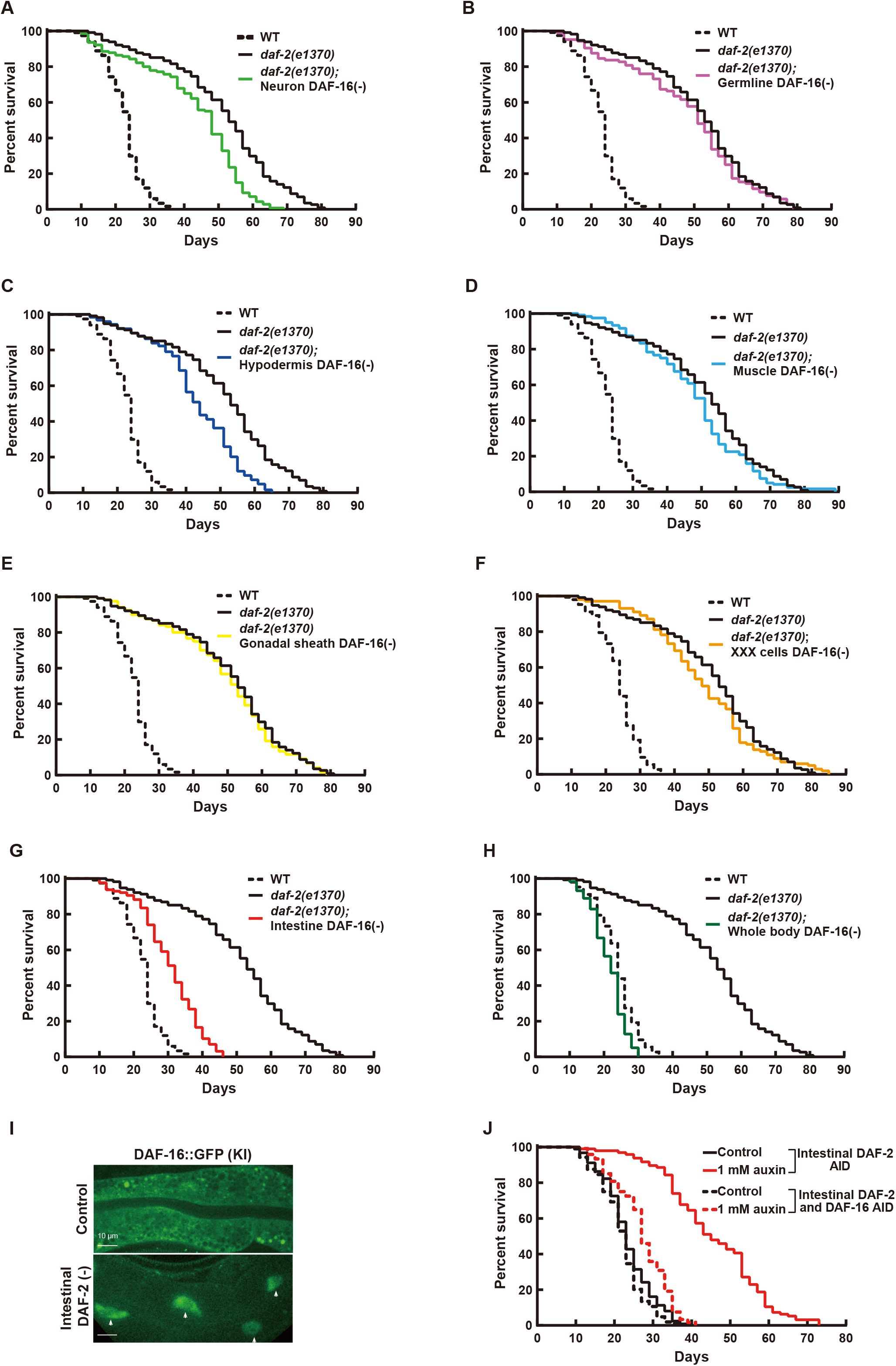
90.3% of the lifespan extension induced by *daf-2(e1370)* required intestinal DAF-16. (A) Degrading DAF-16 from the neurons shortened the *daf-2(e1370)* lifespan by 15.6% (*p*<0.0001). (B) Degrading DAF-16 from the germline had no significant effect on the *daf-2(e1370)* lifespan (*p*>0.05). (C) Degrading DAF-16 from the hypodermis shortened the *daf-2(e1370)* lifespan by 15.5% (*p*<0.0001). (D-F) Degrading DAF-16 in the BWM (D), gonadal sheath (E), or XXX cells (F) had no effect on the lifespan of *daf-2(e1370)* (*p*>0.05). (G) Degrading intestinal DAF-16 shortened the *daf-2(e1370)* lifespan by 40.1% (*p*<0.0001). (H) Degrading DAF-16 in all tissues shortened the *daf-2(e1370)* lifespan by 57.8% (*p*<0.0001). (I) Endogenously expressed DAF-16::GFP accumulated in the intestinal nuclei upon degrading DAF-2 from the intestine. (J) Degrading intestinal DAF-16 largely suppressed the longevity induced by degrading DAF-2 from the intestine (red dashed line versus red solid line, *p*<0.0001). *p*-values are calculated by log-rank tests. See also Table S1 and Figure S5.

Further supporting the above conclusion, we found that endogenously expressed DAF-16::GFP from a KI allele (Figure 2) accumulated in the intestinal nuclei after intestinal DAF-2 was degraded (Figure 4I). Nuclear accumulation is a sign of DAF-16 activation, due to alleviation of the inhibitory phosphorylation by AKT-1/2 on DAF-16 following inactivation of AKT kinase or the upstream kinases AGE-1 and DAF-2 (Kimura et al., 1997; Lin et al., 1997; Ogg et al., 1997; Paradis and Ruvkun, 1998; Pierce et al., 2001). In adult animals, nuclear accumulation of DAF-16::GFP was seen only in the intestine following degradation of DAF-2 in the same tissue. Moreover, 72.7% of the extra lifespan gained by degrading intestinal DAF-2 required intestinal DAF-16 (Figure 4J). Taken together, these results demonstrate that 90% of the lifespan extension from reducing IIS originates in the intestine, and it requires activation of DAF-16 in the same cells.

### 5. Intestine-specific degradation of insulin/IGF-1 receptor extends lifespan without developmental or reproductive defects

Having established that the intestine is the control center for lifespan regulation by IIS, we asked whether lifespan extension may occur without causing the other pleiotropic phenotypes associated with traditional *daf-2* alleles (Gems et al., 1998). We found that intestine-specific degradation of DAF-2, which nearly doubled lifespan (Figure 3G and Figure 4G) caused no developmental abnormality, nor reproductive defects (Figure 5).

**Figure 5.**
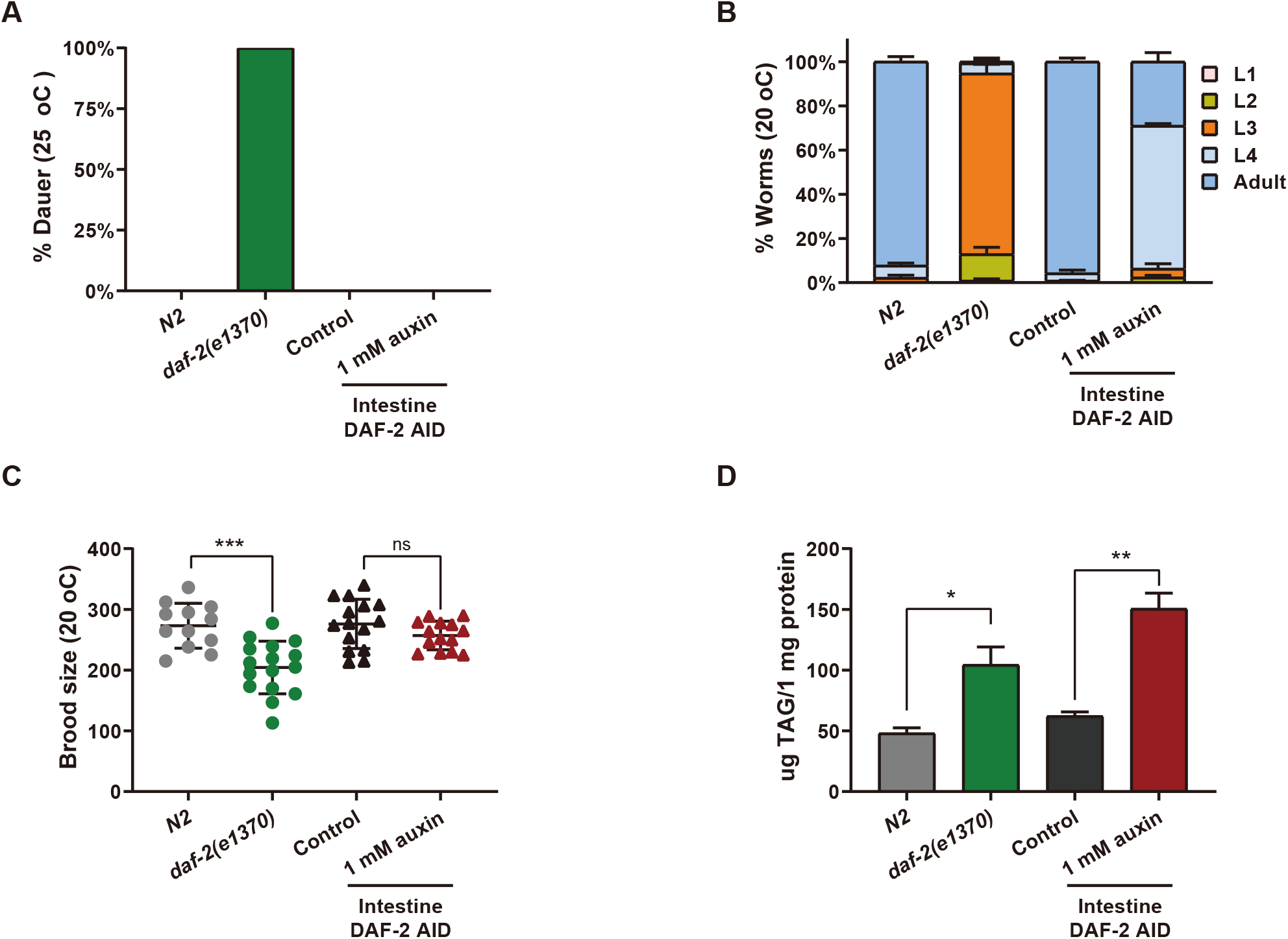
Degrading DAF-2 in the intestine hardly affected development and reproduction. (A) Intestinal degradation of DAF-2 did not cause dauer arrest at 25 °C. (B) The intestinal DAF-2 AID worms developed faster than the *daf-2(e1370)* worms. Data are represented as mean ± SEM. (C) The brood size of intestinal DAF-2 AID worms is comparable to that of the control animals. (D) Degradation of intestinal DAF-2 elevates the triacylglycerol (TAG) content by 2.4-fold relative to the control animals. Data are represented as mean ± SEM. *p*-values are calculated by Student’s *t*-test. ***, *p* < 0.001; **, *p* < 0.01; *, *p* < 0.05; ns, *p* > 0.05.

Compared to the canonical *daf-2(e1370)* allele—whose remarkable longevity is accompanied by mild pleiotropic phenotypes, including 100% dauer formation at 25 °C or ∼0.5% at 20 °C (Gems et al., 1998)—intestinal degradation of DAF-2 caused 0% dauer formation at 25 °C (Figure 5A). The developmental rate of the *daf-2(e1370)* mutant was also significantly slower than WT. A total of 92.3% of freshly laid WT eggs developed into adults after 64 hours at 20 °C, while 81.8% of the *daf-2(e1370)* population were still at the L3 stage. With 93.8% of the population reaching L4 or adulthood under the same conditions, the intestinal DAF-2 AID worms developed faster than the *daf-2(e1370)* worms and slightly slower than WT (Figure 5B).

With respect to brood size, *daf-2(e1370)* animals laid an average of 203 eggs per worm at 20 °C, which is 25% fewer than WT. In contrast, there was no difference in the number of eggs laid per worm between the intestinal DAF-2 AID worms and control worms (Figure 5C).

For the lipid storage phenotype, we found that degrading intestinal DAF-2 elevated the triacylglycerol (TAG) content by 2.4-fold relative to the control animals, which recapitulated the metabolic phenotype of the *daf-2(e1370)* mutant (Figure 5D). Among all the pleiotropic phenotypes examined, this lipid storage phenotype is the only one that remains associated with the long-lived intestinal DAF-2 AID worms. In short, by degrading DAF-2 only in the intestine, the longevity effect was successfully separated from undesirable developmental and reproductive effects associated with a general reduction of IIS.

### 6. The long-lived animals lacking the insulin/IGF-1 receptor in the intestine redefine the molecular signature of longevity

A multitude of downstream effects of IIS means numerous affected genes. A study of 75 publicly available microarray datasets comparing *daf-2(-)* versus *daf-2(-); daf-16(-)* has identified 3,396 DAF-16 target genes, of which 1,663 are up-regulated (Class I targets) and 1,733 are down-regulated (Class II targets) when DAF-16 is active (Tepper et al., 2013). Our next-generation RNA-seq analysis of *daf-2(e1370)* versus WT (N2) worms uncovered 2,463 differentially expressed genes (DEGs) (*p*. adjust<0.05), with 1,509 up-regulated and 953 down-regulated in *daf-2(e1370)* worms (Figure 6A, left panel). Contrasting the KEGG pathways enriched from the Class I or Class II targets (Figure 6A, middle panel) and those from the RNA-seq data, it is evident that RNA-seq data revealed extensive down-regulation of multiple pathways related to metabolism of proteins and RNA (Figure 6A, right panel). Decreased protein metabolism in long-lived *daf-2* worms has been found repeatedly in previous proteomics studies and was shown to contribute positively to the longevity of *daf-2* worms (Stout et al., 2013). Here, we show that this information is in the transcriptome data, discernable using gene set enrichment analysis (GSEA) (Figure 6A, right panel).

**Figure 6.**
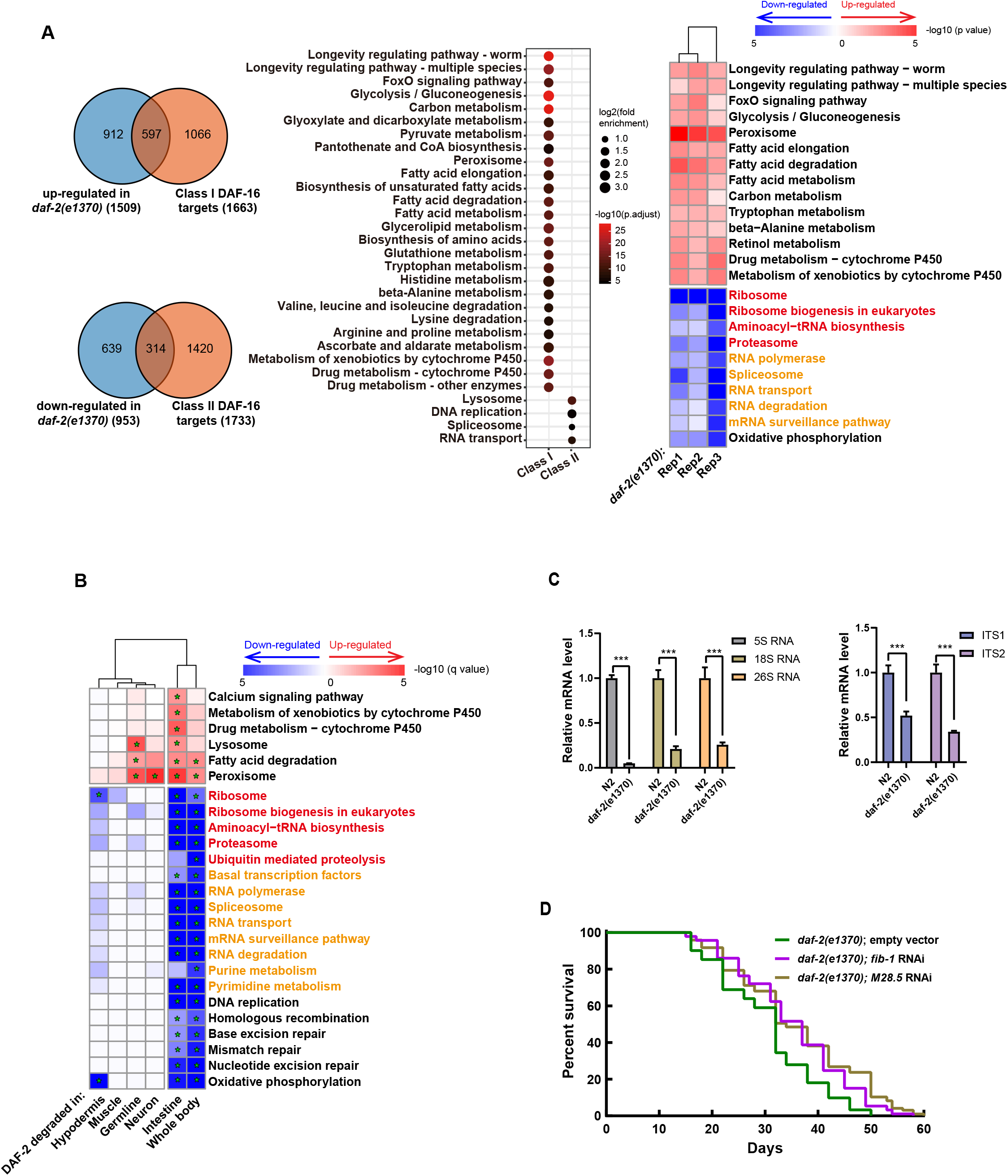
Transcriptional changes induced by degrading intestinal DAF-2 featured down-regulation of RNA and protein metabolism. (A) Transcriptome analysis of *daf-2(e1370)* versus WT (N2) worms. Left panel, overlap of the canonical DAF-16 targets and DEGs (*q*-value <0.05) of *daf-2(e1370)*. Middle panel, enriched KEGG pathways among Class I and Class II DAF-16 targets by Over Representation Analysis (ORA). Only pathways with adjusted *p*-value<0.001 are shown. Right panel, GSEA of *daf-2(e1370)* worms. Pathways with *q*-value<0.01 are shown, and those related to protein metabolism (red color) and RNA metabolism (orange color) are highlighted. (B) GSEA analysis of worms subjected to tissue-specific degradation of DAF-2. Pathways with *q*-value<0.01 in at least one sample are shown, and those related to protein metabolism (red color) and RNA metabolism (orange color) are highlighted. (C) qRT-PCR analysis of ribosomal RNAs (left panel) as well as their precursors (right panel) in *daf-2(e1370)* and N2 worms. Data are represented as mean ± SD. (D) Knocking down of genes related to RNA metabolism (*fib-1* and *M28*.*5*) by RNAi further extended the lifespan of *daf-2(e1370)* worms. See also Table S1, Figure S6 and S7.

We reasoned that with its “clean” longevity phenotype, intestine-specific DAF-2 degradation could help separate gene expression changes associated with longevity from those associated with the developmental or reproductive phenotypes of *daf-2* worms. We thus conducted RNA-seq analysis of worms subjected to tissue-specific degradation of DAF-2. The RNA-seq data of three biological replicates (Figure S6A) showed that the gene expression changes induced by degrading DAF-2 in the intestine and degrading DAF-2 in the whole body are more similar to each other than either is to the other treatments (Figure 6B and Figure S6B-S6F). For both, the up-regulated genes were enriched in fatty acid degradation and peroxisome, and the down-regulated genes were enriched in pathways related to protein metabolism, RNA metabolism, and DNA repair (Figure 6B). “Longevity regulating pathway-worm” was also enriched from the up-regulated genes in the intestinal DAF-2 AID worms at a *q* value <0.05 (Figure S7A).

Notably, RNA-seq analysis underscored down-regulation of genes functioning in protein and RNA metabolism as common features shared by *daf-2(e1370)*, whole-body, and intestine-specific DAF-2 AID (Figure 6A and 6B). Previous studies have shown that decreasing protein synthesis extends lifespan (Depuydt et al., 2013; Lan et al., 2019; Pan et al., 2007; Syntichaki et al., 2007; Tiku et al., 2017) and accounts for ∼40% of the lifespan extension by *daf-2(e1370)* (Li et al., 2021). In comparison, little is known about how down-regulation of RNA metabolism contributes to *daf-2* longevity.

Since ribosomal RNAs constitute the majority of total cellular RNA, we quantified the rRNAs using qRT-PCR. We found that in the *daf-2* mutant, 5S, 18S, and 26S rRNAs all decreased to less than 26% of the WT level (Figure 6C, left panel), while ITS1 and ITS2, which represents the precursor rRNA (pre-rRNA), also decreased to 52% and 34%, respectively (Figure 6C, right panel). Knocking down *fib-1*, which encodes the *C. elegans* fibrillarin, an enzyme that catalyzes 2’-O-methylation of pre-rRNA and U6 snRNA (Hasler et al., 2020; Iyer-Bierhoff et al., 2018), further extended the lifespan of *daf-2(e1370)* worms (Figure 6D). Knocking down *M28*.*5/snu-13*, which encodes another conserved protein functioning in rRNA processing and mRNA splicing (Marmier-Gourrier et al., 2003), also significantly extended the *daf-2* lifespan. Although knocking down *M28*.*5/snu-13* did not affect WT lifespan (Figure S7B), knocking down *fib-1* in WT animals resulted in a longevity phenotype (Tiku et al., 2017; Tiku et al., 2018). These data suggest that a reduction in RNA metabolism contributes positively to *daf-2* longevity.

In summary, intestine-specific degradation of DAF-2 helps identify core gene expression changes underlying the longevity phenotypes of *daf-2* worms, including up-regulation of fatty acid metabolism and peroxisome, and down-regulation of RNA and protein metabolism. These are shared molecular signatures of longevity following different ways of IIS reduction.

### 7. Cross-tissue effect of intestinal insulin/IGF-1 signaling involves non-intestinal FoxO

The finding that degrading DAF-2 specifically in the intestine nearly doubled *C. elegans* lifespan raised another question, namely, whether and how intestinal IIS affects other tissues. To address this question, we constructed tissue-specific GFP reporters in the intestinal DAF-2 AID strain, with which we isolated the cells of interest from day 1 adults either treated with auxin or untreated, for tissue-specific RNA-seq (Figure 7A, Figure S8A and S8B). Principal component analysis (PCA) of the RNA-seq data showed clear distinctions between tissues and between treatments (Figure S8C).

**Figure 7.**
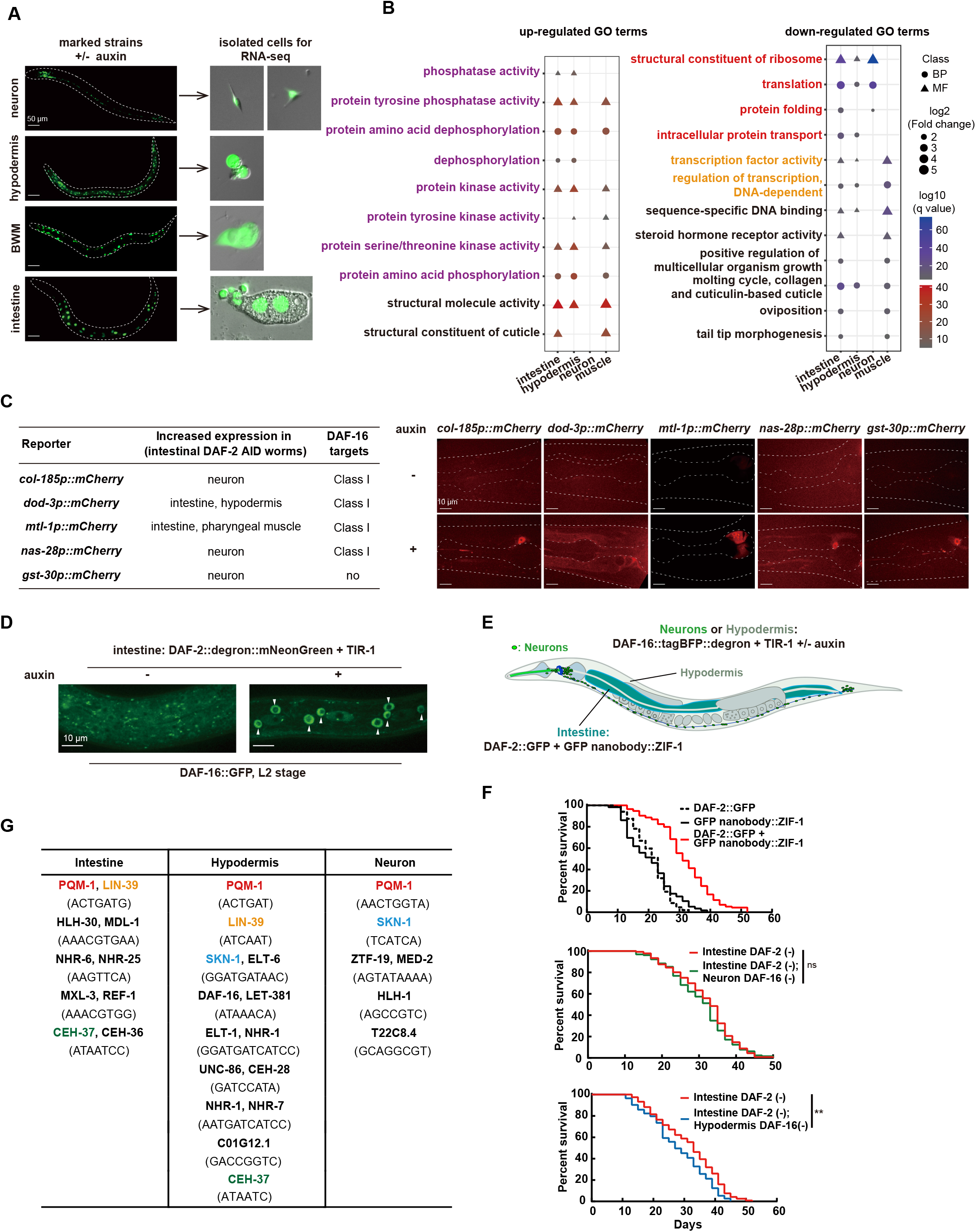
Loss of intestinal DAF-2 triggered gene expression changes in other tissues through cross-tissue DAF-2 to DAF-16 signaling, in part. (A) Isolation of neurons, hypodermis, BWM, and intestine from intestinal DAF-2 AID worms for RNA-seq. Left panel, tissue-specific transgenic reporters in the intestinal DAF-2 AID strain used to isolate tissues of interest. Neurons, labeled by *rgef-1p::NuGFP*; hypodermis, labeled by *dpy-7p::NLS::GFP*; muscle, labeled by *myo-3p::NuGFP*; intestine, labeled by *ges-1p:: NuGFP*. Right panel, representative images of tissue-specific cells isolated by FACS for RNA-seq. (B) GSEA analysis of up-regulated and down-regulated GO terms enriched in each isolated tissue. GO terms with *q*-value<0.01 in at least two tissues are shown. (C) Five DEGs selected to verify the cross-tissue effect of intestinal IIS on non-intestinal tissues by constructing *mCherry* transgenic reporters in the intestinal DAF-2 AID strain. Representative images of each reporter are shown in the right panel. (D) Degrading DAF-2 from the intestine induces DAF-16 nuclear accumulation in the hypodermis at the L2 larval stage. (E) Schematic of the combination of tissue-specific GFP nanobody-mediated ZIF-1 system and tissue-specific AID system to simultaneously degrade DAF-2 and DAF-16 in two different tissues, the intestine and non-intestinal tissues (neurons or hypodermis). (F) Lifespan phenotypes following simultaneous degradation of intestinal DAF-2 and non-intestinal DAF-16. Degrading intestinal DAF-2 by GFP nanobody-mediated ZIF-1 system extended lifespan by 49.5% (top panel). Degrading DAF-16 in the hypodermis (bottom panel), but not in the neurons (middle panel), moderately but significantly decreased the lifespan of the worms in which the intestinal DAF-2 level was reduced. (G) Motif enrichment analysis of transcriptional binding sites among 1-kb promoter sequence of the intestinal, hypodermal, and neuronal DEGs. See also Table S1, Figure S8.

From the isolated intestinal, hypodermal, neuronal, and BWM cells, we found 508, 212, 209, and 26 DEGs, respectively, as a result of degrading DAF-2 in the intestine (Figure S8D). These data demonstrate a clear cross-tissue effect of intestinal IIS on hypodermis and neurons. That only 26 DEGs were found in muscle cells echoes the finding that IIS in muscles is insignificant in lifespan regulation.

Gene ontology (GO) analysis of DEGs of different tissues revealed an intriguing pattern (Figure 7B). Among the ten up-regulated genes that responded to degradation of intestinal DAF-2, eight of the enriched GO terms seen in at least two tissue types are related to protein phosphorylation. GO terms enriched from the down-regulated genes are related predominantly to protein and RNA metabolism, which re-confirms the finding from whole worm RNA-seq data (Figure 6A, right panel, Figure 6B). The intestine displayed a higher degree of similarity with the hypodermis than with the neurons, but down-regulation of structural constituents of ribosome and translation is a shared transcriptional response among all three tissues (Figure 7B). Therefore, decreased protein synthesis, an important mechanism for IIS longevity (Li et al., 2021), is propagated from the intestine to other tissues. Decreased protein synthesis is also a molecular signature of longevity by Dietary Restriction (DR) (Kapahi, 2010; Lan et al., 2019; Stout et al., 2013).

Moderate enrichments of Class I and Class II DAF-16 targets were found among hypodermal and neuronal DEGs (Figure S8E), suggesting that DAF-16 may be activated to some degree in these strains in the absence of intestinal DAF-2. Supporting this idea, four Class I DAF-16 targets were induced outside the intestine following degradation of intestinal DAF-2 (Figure 7C, Figure S8G and S8H). In addition, intestinal DAF-2 degradation induced nuclear accumulation of DAF-16::GFP in the hypodermis during the L1-L2 stage (Figure 7D and Figure S8F).

We then asked whether lifespan extension by intestinal DAF-2 degradation requires DAF-16. We knocked in a tagBFP::degron tag to the genomic locus of *daf-16*, thereby subjecting the expressed tagBFP::degron fusion protein to auxin-induced tissue-specific degradation (Figure 7E). To degrade intestinal DAF-2 in the same animals, we knocked in a GFP sequence to the *daf-2* locus and expressed a GFP nanobody::ZIF-1 fusion protein under an intestinal promoter (Wang et al., 2017). Unlike the AID system, GFP-mediated degradation of DAF-2 was incomplete, but enough to extend lifespan by 49.5% (Figure 7F, top panel). We found that degrading DAF-16 in the hypodermis, but not in the neurons, moderately but significantly shortened the lifespan of worms in which intestinal DAF-2 level was reduced (Figure 7F, middle and bottom panel). The DAF-16 binding site was also enriched among the hypodermal DEGs following the removal of intestinal DAF-2 (Figure 7G). Therefore, we conclude that hypodermal DAF-16 is activated by the loss of intestinal DAF-2 and contributes to longevity.

From the tissue-specific RNA-seq data, we also found evidence suggesting the involvement of other TFs. The PQM-1 and SKN-1 binding sites were significantly enriched in 1-kb promoter sequence of the hypodermal and neuronal DEGs (Figure 7G). These two TFs are required for *daf-2* longevity (Tepper et al., 2013; Tullet et al., 2008). Induction of *gst-30*, which is not a DAF-16 target, was seen in a head neuron after intestinal DAF-2 was degraded (Figure 7C).

Taken together, the above data demonstrate that loss of intestinal DAF-2 induces gene expression changes in other tissues and that there is cross-tissue DAF-2 to DAF-16 signaling.

## Discussion

In contrast to the diverse pleiotropic phenotypes of *daf-2* mutants, previous studies reported restricted and differing expression patterns of *daf-2*, either in neurons and XXX cells based on immunostaining (Kimura et al., 2011) or in the germline based on *in situ* hybridization (Honnen et al., 2012). In this study, we found that in addition to neurons, XXX cells, germ cells, and vulval cells—which display the brightest DAF-2::mNeonGreen signal on the cell surface—the *daf-2* gene is also expressed in the intestine, hypodermis, muscles, and gonadal sheath. We believe that this expression pattern is accurate and complete for the following reasons. First, we engineered the genomic loci of the *daf-2* and *daf-16* genes to label their endogenous protein products with fluorescent tags (Figure1 and Figure 2), thereby avoiding artifacts associated with transgene reporters. Because all the DAF-2 isoforms share the same C-termini, and so do all the DAF-16 isoforms, the C-terminal mNeonGreen or GFP tag should label all DAF-2 or DAF-16 proteins. Second, we developed an ultra-sensitive tagging method (Figure 1A), with which we found that most or all *C. elegans* cells express *daf-2* mRNA (Figure 1C). This ubiquitous expression pattern of *daf-2* overlaps perfectly with that of its downstream TF *daf-16*, and is consistent with the many pleiotropic phenotypes of *daf-2* mutants. For example, mutations of *daf-2* increase lipid storage in the intestine, promote dauer formation during development (which involves the hypodermis producing a dauer cuticle), and suppress induction of multiple vulvae (Nakdimon et al., 2012). Prior to this study, *daf-2* expression had not been detected in the intestine, hypodermis, and vulval cells, where the above phenotypes are found.

Anti-aging by reducing neuronal IIS has been an attractive idea since expression of a *daf-2* transgene under a neuronal promoter, but not a muscle or intestinal promoter, was found to suppress the longevity phenotypes of the *daf-2* mutant (Wolkow et al., 2000). However, transgenic expression of *daf-16* under an intestinal promoter, but not a neuronal or muscle promoter, extended the lifespan of the *daf-2; daf-16* double mutant (Libina et al., 2003). The fact that *daf-2* RNAi readily extends lifespan (Dillin et al., 2002) also casts doubt on this idea because *C. elegans* neurons are refractory to RNAi (Fraser et al., 2000; Kamath et al., 2001). Recently, it was reported that neuronal and intestinal *daf-2* knockout both extend WT lifespan by ∼50% (Uno et al., 2021), as did neuronal and intestinal degradation of DAF-2 (BioRxiv, 2021). A third study showed that intestine-specific DAF-16 degradation shorted the lifespan of *daf-2(e1370)* mutant (Aghayeva et al., 2021). Here, we showed that intestine or neuron-specific degradation of DAF-2 extended lifespan by 94.3% or 18.6%, respectively. Hence, a consensus is forming that intestinal DAF-2 has a substantial effect on lifespan regulation but there is no consensus yet regarding neuronal DAF-2. To solve this controversy, each of the above experiments must be examined to ensure the absence or reduction of neuronal DAF-2 and the intactness of intestinal DAF-2. None of the above studies showed direct evidence of the latter, and only this study showed direct evidence of the former. Our study is the only one so far that determined the expression pattern of *daf-2*, and the DAF-2::mNeonGreen or *daf-2::degron::mNeonGreen* KI strain should be a useful tool to visualize the effect of *daf-2* knockdown, knock out, or DAF-2 protein degradation in neurons. Pertaining to the intactness of intestinal DAF-2 in our degradation experiments of neuronal DAF-2, although the weak intestinal DAF-2::mNeonGreen signal did not provide a direct readout, we reason that intestinal DAF-2 was not affected when neuronal DAF-2 was degraded based on the following: (1) neuronal DAF-2 was verifiably degraded; (2) this treatment extended lifespan by 18.6%; (3) intestine-specific degradation of DAF-2 extended lifespan by 94.3%; and (4) if the 18.6% lifespan extension had resulted from an unintended loss of intestinal DAF-2, it would further refute the idea of IIS regulating lifespan from neurons.

Related to this, mechanistic studies of how certain neuronal changes extend lifespan, *e*.*g*. the mitochondrial unfolded protein response (UPR^mt^), expression of XBP-1s, and activating neuronal CRTC-1, often identify the intestine as the responsive organ (Burkewitz et al., 2015; Durieux et al., 2011; Imanikia et al., 2019; Rera et al., 2013; Zhang et al., 2018b). Many signaling pathways that influence lifespan converge on DAF-16, including IIS, AMPK signaling, mitochondrial signaling, CRCT-1, and germline signaling (Burkewitz et al., 2015; Kenyon, 2010). Activation of intestinal DAF-16, in particular, underlies a growing number of lifespan-extending conditions, including germline ablation (Berman and Kenyon, 2006; Libina et al., 2003; Lin et al., 2001), overexpression of HSF-1 in neurons (Douglas et al., 2015), temperature-dependent lifespan regulation by IL1 and ASJ neurons (Zhang et al., 2018a), and the classic case of lowering IIS (Libina et al., 2003; Uno et al., 2021); this study). Lifespan extension by activating intestinal DAF-16 is not unique to *C. elegans*. The worm intestine, in addition to being a digestive tract, doubles as an adipose tissue. In mice, adipose tissue-specific knockout of insulin receptor (FIRKO mice) extends lifespan (Bluher et al., 2003), and intestinal epithelium-specific knockout of IR alleviates insulin resistance in aged animals (Ussar et al., 2017). Both treatments activate the mammalian counterpart of DAF-16, the FoxO TFs, in the IR KO cells. It would be interesting to find out whether intestine-epithelium-specific knock-out of IR extends lifespan in the mouse model.

Based on the findings in mice described above and the finding in *C. elegans* that depletion of intestinal DAF-2 doubles lifespan without adverse developmental or reproductive effects (this study and Venz et al., 2021, BioRxiv), we suggest that intestine-specific reduction of IIS is a potentially promising approach to fight against aging in mammals.

The intestine supplies nutrients to the entire body, making it plausible that intestinal IIS may regulate organismal aging by altering nutrient supply. IIS is a master regulator of glucose metabolism and lipid metabolism across the animal kingdom (Chatterjee and Perrimon, 2021; Zhang and Liu, 2014). In *C. elegans*, IIS also positively regulates protein synthesis (Li et al., 2021; Stout et al., 2013). The transcriptional changes shared by whole-body and intestine-specific DAF-2 degradation include up-regulation of peroxisome and fatty acid degradation, and down-regulation of RNA and protein metabolism. We speculate that these altered metabolic pathways subsequently alter the export of nutrients such as the building blocks of lipids, RNAs, and proteins out of the intestine to the other tissues.

In light of our findings, the genetic regulation of lifespan by IIS and the dietary regulation of lifespan are unified in the intestine, in nutrition and metabolism, and in down-regulation of protein synthesis. These likely represent the nexus of additional anti-aging strategies because the intestine is frequently found to be the responsive tissue or organ downstream of many lifespan-extending conditions, as discussed above. We therefore think that these findings will help focus the ever-expanding directions of aging research.

## Methods

### *Carenorhabdities elegans* Maintenance

Nematodes were maintained at 20 °C on standard nematode growth medium (NGM) agar plates seeded with *Escherichia coli* OP50 unless otherwise stated (Brenner, 1974).

#### Strain construction

##### 1) Transgenic strains

To generate transgenic animals carrying extrachromosomal arrays (hqEx), 5-50 ng/µl of the indicated plasmid was injected to the gonad using standard method.

##### 2) Knocking-in strains

CRISPR engineering for all knocking-ins was performed by microinjection using the homologous recombination approach (Dickinson et al., 2013). The injection mix contained at least two Cas9-sgRNAs expressing plasmids (50 ng/ μl for each), a selection marker pRF4 (*rol-6(su1006)*, with roller phenotype) (50 ng / μl), and a homologous recombination plasmid (50 ng/μl). To generate the sgRNA plasmids, primers were designed with the CRISPR DESIGN tool (https://zlab.bio/guide-design-resources) and inserted into the pDD162 vector (Addgene) using the site-directed mutagenesis kit (TOYOBO). To generate the homologous recombination (HR) plasmids, two homologous arms (∼1,000 bp each) corresponding to the 5′- and 3′-sides of the insertion site, respectively, were cloned into the vector. All injection plasmids were purified with PureLink PCR Micro kit (Invitrogen) and injected into the gonad of young adult hermaphrodite worms using standard method. F1s with roller phenotype were singled on a new NGM plate and allowed to produce sufficient offspring. Successful knock-in events were screened by PCR genotyping from independent F1 transgenic animals’ progeny that did not display roller phenotype, and further confirmed by DNA sequencing.

Strains and sgRNAs used in this study are listed in Table S2 and Table S3, respectively.

#### Confocal imaging

Confocal image of Figure 1D (left panel) was captured by ZEISS LSM 880 microscope equipped with a 63×, 1.4 numerical aperture oil-immersion objective as Z-stacks of 1 μm-thick slices under the lambda-mode (settings: pixel dwell: 2.06 μs; average: line 1; master gain: 750; pinhole size: 91 μm; filter: 411-695 nm; beam splitter: MBS 488; lasers: 488 nm, 30 %). Images were processed using ZEN software (Carl Zeiss Inc.).

All other confocal images were taken using the spinning-disk microscope (UltraVIEW VOX; PerkinElmer) equipped with a 63×, 1.4 numerical aperture oil-immersion objective, except for those in Figure 7A (left panel) and Figure S8 (G-H) taken using 10× objective. Images were viewed and processed using Volocity software (PerkinElmer).

Worms were anaesthetized using 1 mM levamisole hydrochloride in water on 3 % agarose pads on glass slides. In all the imaging studies, images within the same figure panel were taken with the same parameter and adjusted with identical parameters using ImageJ software.

#### Auxin treatment

Auxin treatment was performed by transferring worms to OP50-seeded NGM plates containing 1 mM auxin as previously described (Zhang et al., 2015). Briefly, the 400 mM natural auxin indole-3-acetic acid (IAA) (Alfa Aesar) stock solution in ethanol was prepared freshly. Then, the stock solution was added into the NGM agar cooled to about 50 °C in water bath before pouring plates. For all auxin treatment experiments, 0.25% ethanol was used as a control. 100 ul of OP50 overnight culture was seeded onto the auxin or control NGM agar plates (60 × 15 mm). Aluminum foil was used to protect the auxin-containing plates with from light, and then kept plates at room temperature for 1-2 days before use.

#### Lifespan analysis

All lifespan assays were performed at 20 °C unless otherwise stated. Strains were synchronized by allowing 40 gravid adults to lay eggs for 4 hours on OP50-seeded NGM plates at 20 °C. Then, approximately 150 worms at early adulthood stage were transferred on ten fresh OP50-seeded NGM plates containing 1mM auxin or 0.25% ethanol. Animals were transferred to new plates every day until the end of reproductive period, after which worms were transferred to fresh plates every week. For lifespan assays involved degrading DAF-2 throughout the whole body (Figure 3H), 50 μg/ml 5-fluoro-deoxyuridine (FUdR) was also added into the NGM plates to prevent its internal egg hatching phenotype from interfering with the lifespan measurement. In this experiment, worms were transferred to fresh plates every 4 days until death.

Live worms were scored every 2 days. A worm was considered as death if it showed no response to the gentle touch with a platinum wire on head and tail. Worms that had internally hatched larvae (‘bagged’) or ruptured vulvae (‘exploded’) or crawled off the agar surface or became contaminated were censored from the analysis. Statistical analyses were performed using IBM SPSS Statistics 20 software. *p* values were calculated using the log-rank (Mantel-Cox) method.

#### DAF-16 nuclear translocation

To analyze DAF-16 nuclear localization, worms were cultured at 20 °C on NGM plates containing 0.25% ethanol or 1 mM auxin from eggs laid within a 4-hour period and imaged at the indicated stage in the figure (Figure 4I, day 1 of adulthood; Figure 7D, L2 larval stage; Figure S8F, L1 larval stage). As soon as worms were removed from incubation, they were mounted on slides and imaged immediately.

#### Dauer assay

For the dauer assay of N2 and *daf-2(e1370)* worms, 25-30 gravid adults were allowed to lay eggs for 1 hour on OP50-seeded standard NGM plates at 20 °C. For the dauer assay of intestinal DAF-2 AID worms, eggs were laid on NGM plates containing 0.25% ethanol or 1 mM auxin. After picking off the adults, the resulting synchronous population was transferred to 25 °C and scored dauer formation after 72 hours post hatch.

#### Developmental Assay

We semi-synchronized the worms by allowing 30 gravid adults to lay eggs 2 hours at 20 °C and then removing out the adult worms. After 64 hours, the developmental state of the worms was determined.

#### Brood size assay

16 synchronized L4 stage worms were singled on individual NGM plates seeded with OP50. The animals were transferred to fresh plates every 24 hours for 4-5 days. Worms that crawled off the plates, bagged or exploded were censored from the analysis. The number of hatched worms was counted two days later. The *p* values were determined by Student’s t-test.

#### TAG measurement

TAG measurement was performed as previously described (Li et al., 2017). Briefly, 8000 synchronized L1 larval worms were placed on each 10 cm OP50-seeded NGM plate with or without auxin and collected at late L4 stage. One-eighth of worms was taken out to extract total soluble protein, and the protein concentration was quantified using the BCA Protein Assay Kit (Pierce). The sample for TAG measurement was homogenized and 20 μg tri-C17:0 TAG (Nu-Chek) was added as an internal calibration standard, followed by the extraction of total lipid. TAG was separated from total lipid on a thin-layer chromatography (TLC) plate, then transmethylated with 2 ml methanol and 50 μl sulfuric acid to prepare fatty acid methyl esters (FAMEs). FAMEs were re-dissolved in 2 ml pentane and chromatographed using a GC/MS instrument (SHIMADZU, QP2010 Ultra) with a DB-23 GC column (Agilent, 122-2332). FAME peaks were identified according to the fatty acid standards and integrated to calculate the TAG amount, which was finally normalized by the protein amount.

#### qRT-PCR

Total RNAs were extracted from N2 and *daf-2(e1370)* worms on adult day 1 using TRIzol (INVITROGEN), followed by using DNase I to remove contaminant DNA. The cDNA was synthesized by using a reverse transcription kit (TAKARA). qPCR was carried out on an ABI 7500 Fast real-time PCR system using a TAKARA real-time PCR kit (SYBR Premix Ex TaqTM II). mRNA levels of *pmp-3* and *act-1* were used as the internal control.

#### Preparation worm samples for RNA sequencing at the whole worm level

Six AID worm samples of degrading DAF-2 specifically in the intestine, neuron, hypodermis, muscle, germline, or whole body and one control worm (MQD2428) were prepared for RNA sequencing with three biological replicates. Synchronized L1 worms were initially cultured on the standard high-growth (HG) plates supplemented with OP50 for 24 hours at 20 °C to prevent entering the dauer stage, and then transferred onto HG plates containing 1 mM auxin to degrade the DAF-2 protein in different tissues. Worms were washed with M9 buffer and harvested on adult day 1.

#### Isolation of tissue-specific cells from the intestine-specific DAF-2 AID worms by FACS

Synchronized day 1 adult transgenic worms with GFP-labeled neurons, muscle, hypodermis, or intestine (*rgef-1p::NuGFP, myo-3p::NuGFP, dpy-7p::NLS::GFP*, and *ges-1p::NuGFP*) were prepared for cell isolation. Cells were isolated following the procedure described in (Kaletsky et al., 2016; Kaletsky et al., 2018) with minor modifications. Briefly, worms were briefly subjected to SDS-DTT treatment, proteolysis, and mechanical disruption. Cell suspensions were gently passed over a 5 μm syringe filter (Millipore) for neuron cell isolation; 20 μm filter (PluriSelect) for muscle and hypodermal cell isolation. To isolate intestinal cells, cell suspensions were passed through a 40 μm cell strainer (Falcon) and then spun at 800 x g for 3 min in a tabletop centrifuge. The filtered cells were diluted in PBS/20% FBS and sorted using BD FACS Aria II (BD Biosciences). Gates for detection were set by comparison to MQD2428 (hqKi363[*daf-2::mNeonGreen*]) cell suspensions prepared on the same day from a population of worms synchronized alongside the experimental samples. Positive fluorescent events were sorted directly into tubes containing Trizol LS (Invitrogen) for subsequent RNA extraction.

#### RNA sequencing and data pre-processing

RNAs were isolated using a standard Trizol/chloroform/isopropanol method, RNA quality and quantity were assessed using the Agilent 2100 Bioanalyzer RNA Pico chip (Thermo Fisher Scientific). For whole worm RNA-seq, libraries were prepared using the NEBNext Ultra RNA library Prep Kit (NEB) according to the manufacturer’s instructions, and then sequenced using an Illumina HiSeq X Ten System in the paired-end mode (2 × 150 bp) through the service provided by Bionova. To construct RNA-seq libraries of FACS isolated tissue-specific cell samples, RNAs (200 ∼500 pg) were amplified with oligo-dT, then reverse transcribed to cDNA based on polyA tail. The template was switched to the 5’ end of the RNA and the full-length cDNA was generated by PCR. Purified cDNA was fragmented into small pieces with fragment buffer by PCR, and the product was purified and selected by the Agencourt AMPure XP-Medium kit (Thermo Fisher Scientific). cDNA was quantified by Agilent Technologies 2100 bioanalyzer. The double stranded PCR product undergo QC step was heat denatured and circularized by the splint oligo sequence. The single strand circle DNA (ssCir DNA) was formatted as the final library. The final library was amplified with phi29 (Thermo Fisher Scientific) to make DNA nanoball (DNB) which had more than 300 copies of one molecular, DNBs were loaded into the patterned nanoarray and single end 100 bases reads were generated on BGISEQ500 platform (BGI-Shenzhen, China).

FASTQC (version 0.10.1) was used to inspect the quality of the raw sequencing data. The raw data were filtered to remove primers, contamination, and low-quality reads, then the Illumina adapter sequences were trimmed to obtain the clean sequencing data. The reads were aligned to the *C. elegans* reference genome (wbcel235.97) using HISAT2 (version 2.1.0) (Kim et al., 2015) with Ensembl gene annotations (using default parameters). Mapped reads that overlap with coding gene features were counted using featureCounts (version 1.6.5) (Liao et al., 2014). The total number of mapped clean reads for each library ranged from 25-35 million.

#### Differential expression (DE) analysis

For the whole worm RNA-seq data, The R package DESeq2 (version 1.24.0) (Love et al., 2014) was used for data normalization and differential expression analysis. Data quality was assessed by hieratical clustering of samples and principal component analysis (PCA). Differentially expressed genes between auxin treatment and control samples were determined using the *deseq* function which is based on the Negative Binomial (Gamma-Poisson) distribution. For the tissue-specific RNA-seq data, the R package edgeR (version 3.26.8) (McCarthy et al., 2012) was used for multidimensional scaling (MDS), PCA and differential expression analysis. Four outliers (rep 5 of neuron cell control sample, rep 1 of intestine cell auxin treatment sample, rep 1 of hypodermis cell control sample, and rep 2 of muscle cell auxin treatment sample) were removed from further analysis. Differentially expressed genes between auxin treatment and control samples were determined using the *glmQLFit* function (using the parameter “robust=TRUE”) which is based on the quasi-likelihood (QL) F-test.

#### Gene Ontology (GO) and KEGG pathway enrichment analysis

The R package gage was used to identify enrichment of GO terms and KEGG pathways which are based on a Generally Applicable Gene-set Enrichment (GAGE) method (version 2.34.0) (Luo et al., 2009). The package clusterProfiler (version 3.12.0; (Yu et al., 2012)) was used to analyze the over-represented GO terms within up- and down-regulated gene. GO term or KEGG pathway with an adjusted *p*-value <0.01 was defined as significantly changed unless otherwise noted. Data visualization was performed using the R package ggplot2 (version 3.3.2; Wickham H (2016)).

#### TF binding motif analysis

1.0 kb promoter regions upstream of the DE genes from tissue-specific RNA-seq data were retrieved online from WormBase using the Parasite Biomart tool. The motif matrices were identified using RSAtools (van Helden et al., 1998), and then analyzed using footprintDB (Sebastian and Contreras-Moreira, 2014) to identify the potential transcription factors predicted to bind to similar DNA motifs.

#### Quantification and statistical analysis

Statistical analyses of lifespan assays were performed via the IBM SPSS Statistics 20 software. Detailed description of tests performed to determine statistical significance is included in figure legends. *, *p* < 0.05; **, *p* < 0.01; ***, *p* < 0.001; ****, *p* < 0.0001; ns, *p* > 0.05.

## Acknowledgments

We thank the Caenorhabditis Genetics Center (CGC), which is supported by the NIH Office of Infrastructure Programs (P40 OD010440), for providing worm strains. We also thank Drs. Guang-Shuo Ou and Xiao Liu for providing us worm strains and plasmids. This work was supported by Ministry of Science and Technology of China (2014CB84980001 to M.-Q.D.), Beijing Municipal Science and Technology Commission (a fund for cultivation and development of innovation base), and National Natural Science Foundation of China (NSFC-ISF 30261143020 to M.-Q.D.).

## Author contributions

Y.-P.Z., W.-H.Z., and M.-Q.D. designed the experiments and interpreted the data. Y.-P.Z. and W.-H.Z. performed most of the experiments. P.Z. performed the qRT-PCR experiments and lifespan assays related to RNA metabolism. Q.L. performed the TAG quantification experiment. Y.S. performed the 3D imaging experiment. J.-W.W. performed some of the bioinformatics analysis. S.-B.Z., T.C., and C.Z. involved in the interpretation of some of the data. Y.-P.Z., W.-H.Z., and M.-Q.D. drafted and revised the manuscript. M.-Q.D. supervised this study.

## Declaration of interests

The authors declare no competing interests.

## Supporting information

**Figure S1. Knocking in *daf-16::gfp* or *daf-2::mNeonGreen* does not affect lifespan or development. Related to Figure 1 and 2**.

(A) The lifespan of *daf-16::gfp* knock-in worms or *daf-2::mNeonGreen* knock-in worms is comparable to that of WT animals.

(B) *daf-16::gfp* does not affect the Daf-c phenotype of *daf-2(e1370)* mutant at 25 °C, and *daf-2::mNeonGreen* knock-in does not induce dauer formation at 25 °C. Data shown are one of three biological replicates.

**Figure S2. The spatiotemporal expression pattern of *daf-2p*::NuGFP. Related to Figure 1**.

(A) DAF-2::mNeonGreen is co-expressed with ciliated sensory neuron maker *osm-6p::mCherry*. (B-G) NuGFP reporter shows the expression pattern of *daf-2* at the two-cell embryonic stage (B), embryonic stage (C), L1 larval stage (D), L2 larval stage (E), L3 larval stage (F), and L4 larval stage (G).

**Figure S3. The spatiotemporal expression pattern of DAF-16::GFP. Related to Figure 2**.

The expression pattern of DAF-16::GFP at embryonic stages (A), L1 larval stage (B), L2 larval stage (C), L3 larval stage (D), and L4 larval stage (E).

**Figure S4. Tissue-specific degradation of DAF-2::degron::mNeonGreen by the AID system. Related to Figure 3**.

(A) Schematic of the AID system. One key component of the AID system is a plant-specific F-box protein called transport inhibitor response 1 (TIR1), which is expressed under the control of a tissue-specific promoter to achieve spatial specificity. The other key component, an auxin-inducible degron sequence from the IAA17 protein, is fused to the protein of interest. Once bound to the plant hormone, auxin, TIR1 targets degron-tagged proteins for ubiquitin-dependent proteasomal degradation. The *C. elegans* AID system employs the *Arabidopsis thaliana* TIR1 and a 44-amino acid minimal degron sequence.

(B) Schematic of construction of tissue-specific AID worms by CRISPR/Cas9 genomic editing. The *eft-3* promoter is replaced with neuron-specific, hypodermis-specific, body wall muscle-specific, or gonadal sheath-specific promoter on chromosome II. The *sun-1* promoter and *sun-1* 3’UTR are replaced with XXX cells-specific promoter and unc-54 3’UTR, respectively, on chromosome IV. (C-J) Specifically degrading DAF-2 in the neurons (C), intestine (D), body wall muscles (E), hypodermis (F), germline (G), gonadal sheath (H), XXX cells (I), or whole body (J) with 1 mM auxin treatment.

**Figure S5. Tissue-specific degradation of DAF-16::GFP::degron in *daf-2(e1370)* worms by the AID system. Related to Figure 4**.

(A-H) Specifically degrading DAF-16::GFP::degron in the neurons (A), intestine (B), body wall muscles (C), hypodermis (D), germline (E), gonadal sheath (F), XXX cells (G), or whole body (H) with 1 mM auxin treatment.

**Figure S6. Transcriptome analysis of tissue-specific DAF-2 AID worms. Related to Figure 6**.

(A) Principal component analysis (PCA) shows that samples cluster with their respective genotypes. R1, R2, and R3 represent three biological replicates.

(B-F) Overlap of DEGs in the whole body DAF-2 AID worms and that in the intestinal DAF-2 AID worms. (B), hypodermal DAF-2 AID worms (C), neuronal DAF-2 AID worms (D), germline DAF-2 AID worms (E), or body wall muscle DAF-2 AID worms (F). DEGs were defined as adjusted *p*-value<0.001.

**Figure S7. KEGG enrichment of tissue-specific DAF-2 AID worms by the GSEA analysis. Related to Figure 6**.

(A) KEGG pathways enriched in each tissue-specific DAF-2 AID worm. Pathways with *q*-value<0.05 are shown.

(B) Knocking down of *M28*.*5* does not affect WT lifespan.

**Figure S8. Tissue-specific transcriptome analysis of the intestinal DAF-2 AID worm. Related to Figure 7**.

(A) Workflow of tissue-specific cell isolation by FACS.

(B) Sample information of isolated cells for tissue-specific RNA-seq.

(C) PCA analysis shows that samples cluster with their respective tissue types as well as corresponding treatment (ethanol versus auxin).

(D) Numbers of DEGs (FDR < 0.05) identified in each isolated tissue in the intestinal DAF-2 AID worms.

(E) Moderate enrichments of Class I and Class II DAF-16 targets are found among the hypodermal and neuronal DEGs upon degrading DAF-2 in the intestine. *p*-values are calculated using a hypergeometric test.

(F) Degrading DAF-2 from the intestine induces DAF-16 nuclear accumulation in the hypodermis at the L1 larval stage.

(G-H) Representative images showing that intestinal DAF-2 degradation regulates the expression of classical Class I DAF-16 targets *mtl-1p::mCherry* (G) and *dod-3p::mCherry* (H) in the same tissue.

**Table S1. Statistical analyses of lifespan experiments (related to Figure 3, 4, 6 and 7)**

**Table S2. Strains used in this study**.

**Table S3. Oligonucleotide sequences used in this study**.

## Notes

### Competing Interest Statement

The authors have declared no competing interest.

