## Supplemental figures and tables for "Degrading intestinal DAF-2 nearly doubles *Caenorhabditis elegans* lifespan without affecting development or reproduction"

#### Figure S1

A

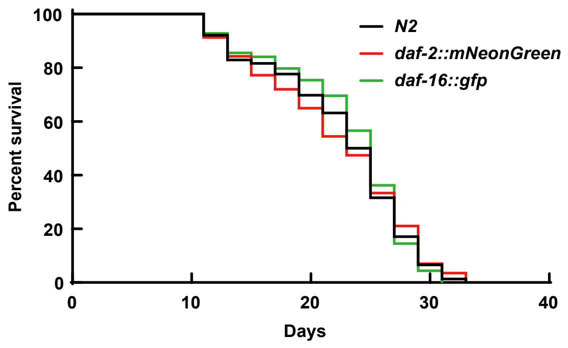

Percent survival
Days
N2
daf-2::mNeonGreen
daf-16::gfp


B

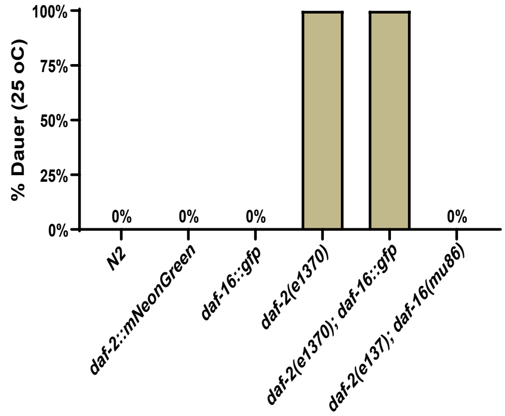

% Dauer (25 oC)
N2
daf-2::mNeonGreen
daf-16::gfp
daf-2(e1370)
daf-2(e1370); daf-16::gfp
daf-2(e137); daf-16(mu86)

### Figure S2

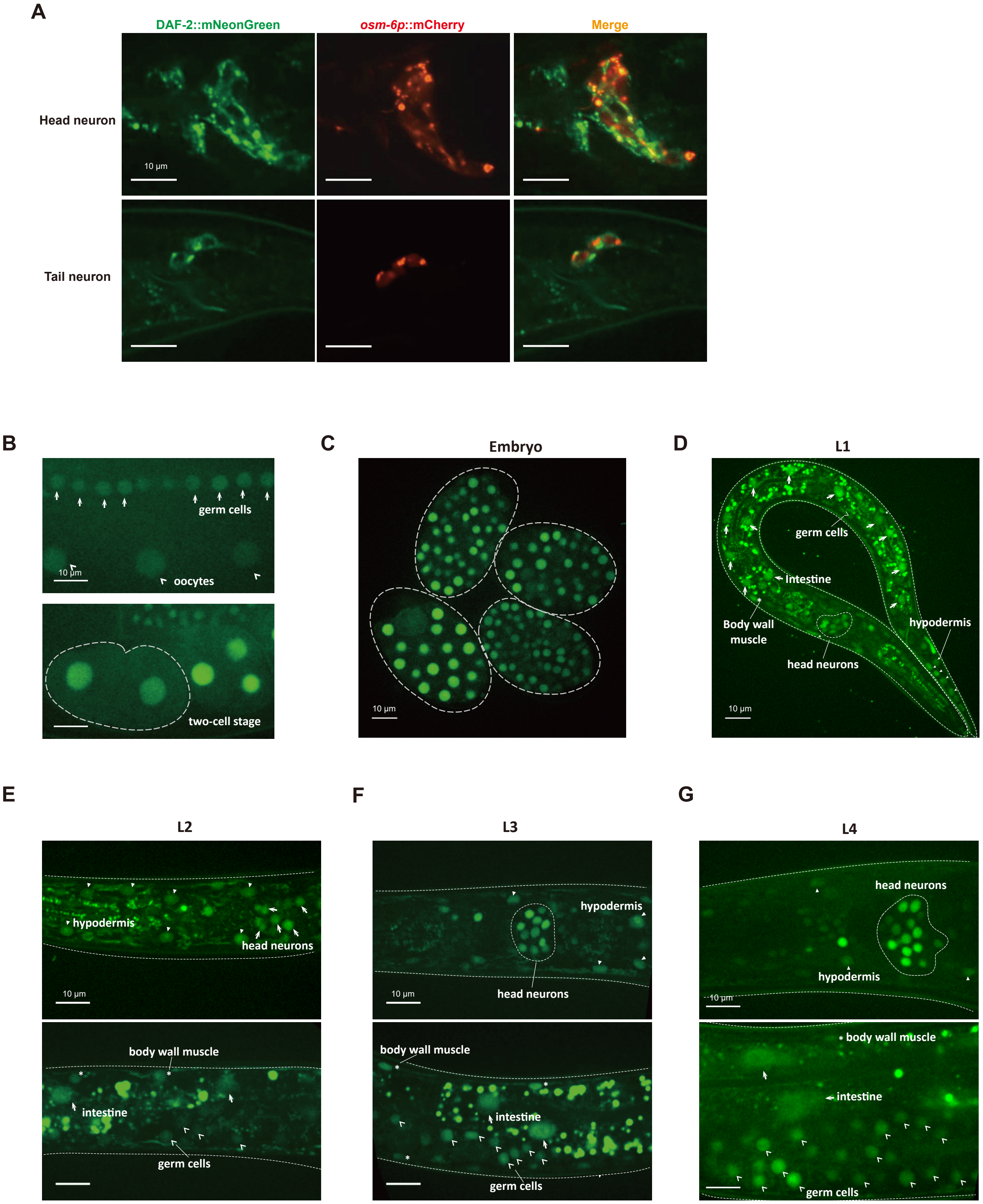
A
DAF-2::mNeonGreen
osm-6p::mCherry
Merge
Head neuron
Tail neuron
10 μm
B
germ cells
10 μm
oocytes
two-cell stage
C
Embryo
10 μm
D
L1
germ cells
intestine
Body wall muscle
hypodermis
head neurons
10 μm
E
L2
hypodermis
head neurons
10 μm
body wall muscle
intestine
germ cells
F
L3
hypodermis
head neurons
10 μm
body wall muscle
intestine
germ cells
G
L4
head neurons
hypodermis
10 μm
body wall muscle
intestine
germ cells

### Figure S3

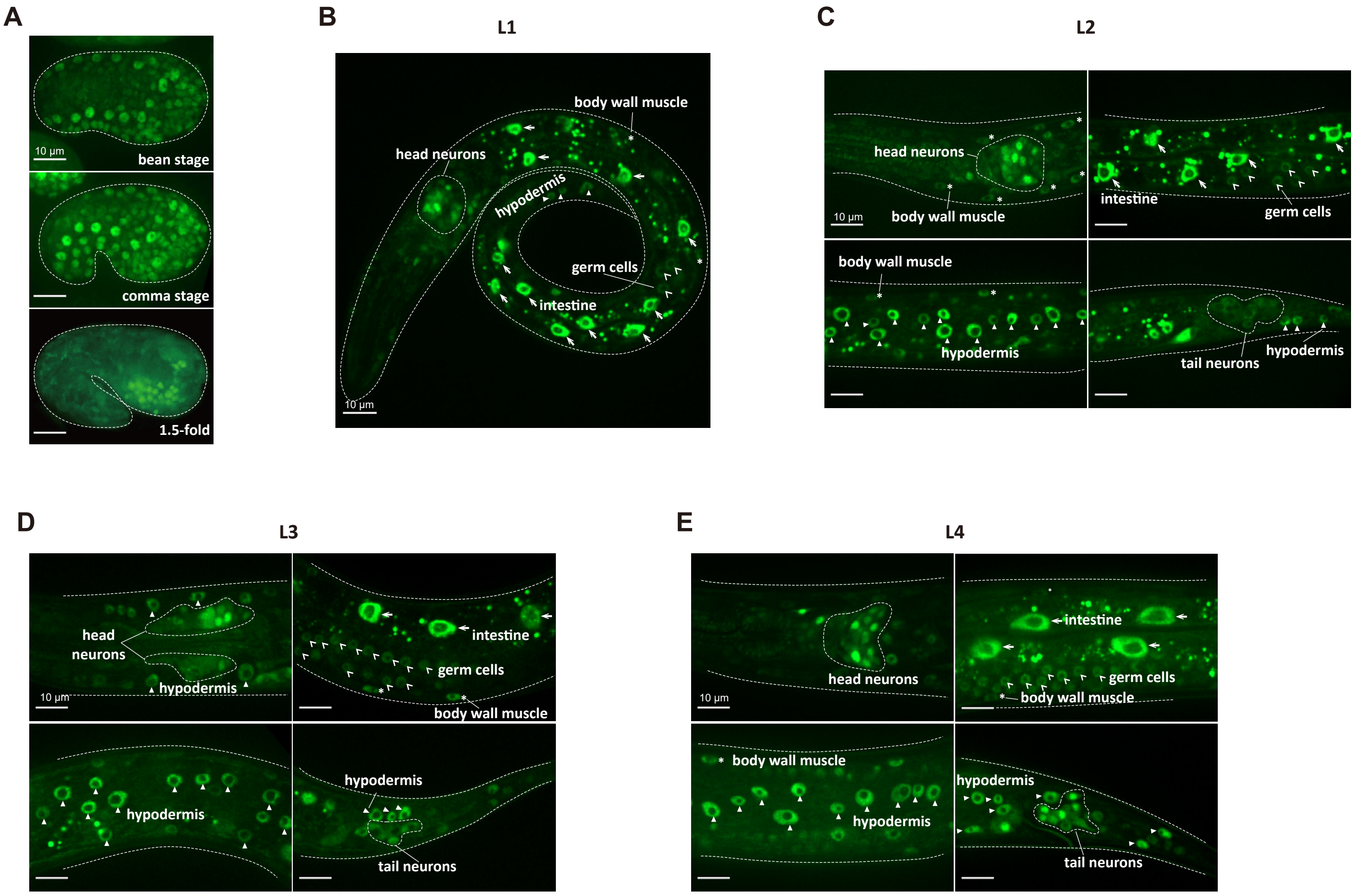

A
bean stage
comma stage
1.5-fold
10 μm
B
L1
head neurons
body wall muscle
hypodermis
germ cells
intestine
10 μm
C
L2
head neurons
body wall muscle
intestine
germ cells
hypodermis
tail neurons
10 μm
D
L3
head neurons
hypodermis
intestine
germ cells
body wall muscle
tail neurons
10 μm
E
L4
head neurons
intestine
germ cells
body wall muscle
hypodermis
tail neurons
10 μm

### Figure S4

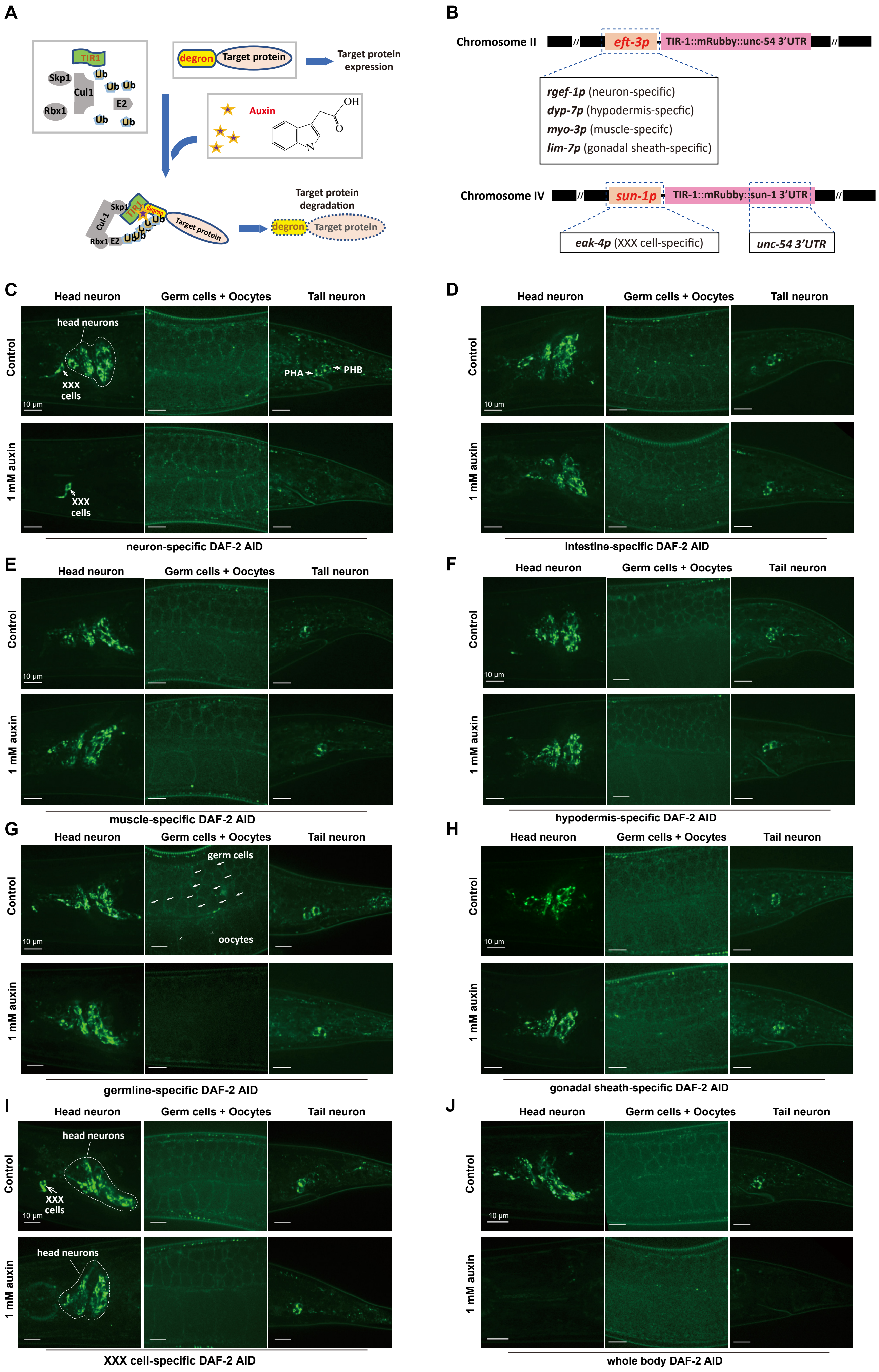

A
TIR1
Skp1
Cul1
Rbx1
Ub
E2
degron
Target protein
Target protein expression
Auxin
OH
O
N
Target protein degradation
B
Chromosome II
eft-3p
TIR-1::mRubby::unc-54 3'UTR
rgef-1p (neuron-specific)
dyp-7p (hypodermis-specfic)
myo-3p (muscle-specifc)
lim-7p (gonadal sheath-specific)
Chromosome IV
sun-1p
TIR-1::mRubby::sun-1 3'UTR
eak-4p (XXX cell-specific)
unc-54 3'UTR
C
Head neuron
Germ cells + Oocytes
Tail neuron
Control
1 mM auxin
head neurons
XXX cells
PHA
PHB
10 μm
neuron-specific DAF-2 AID
D
intestine-specific DAF-2 AID
E
muscle-specific DAF-2 AID
F
hypodermis-specific DAF-2 AID
G
germ cells
oocytes
germline-specific DAF-2 AID
H
gonadal sheath-specific DAF-2 AID
I
XXX cell-specific DAF-2 AID
J
whole body DAF-2 AID

### Figure S5

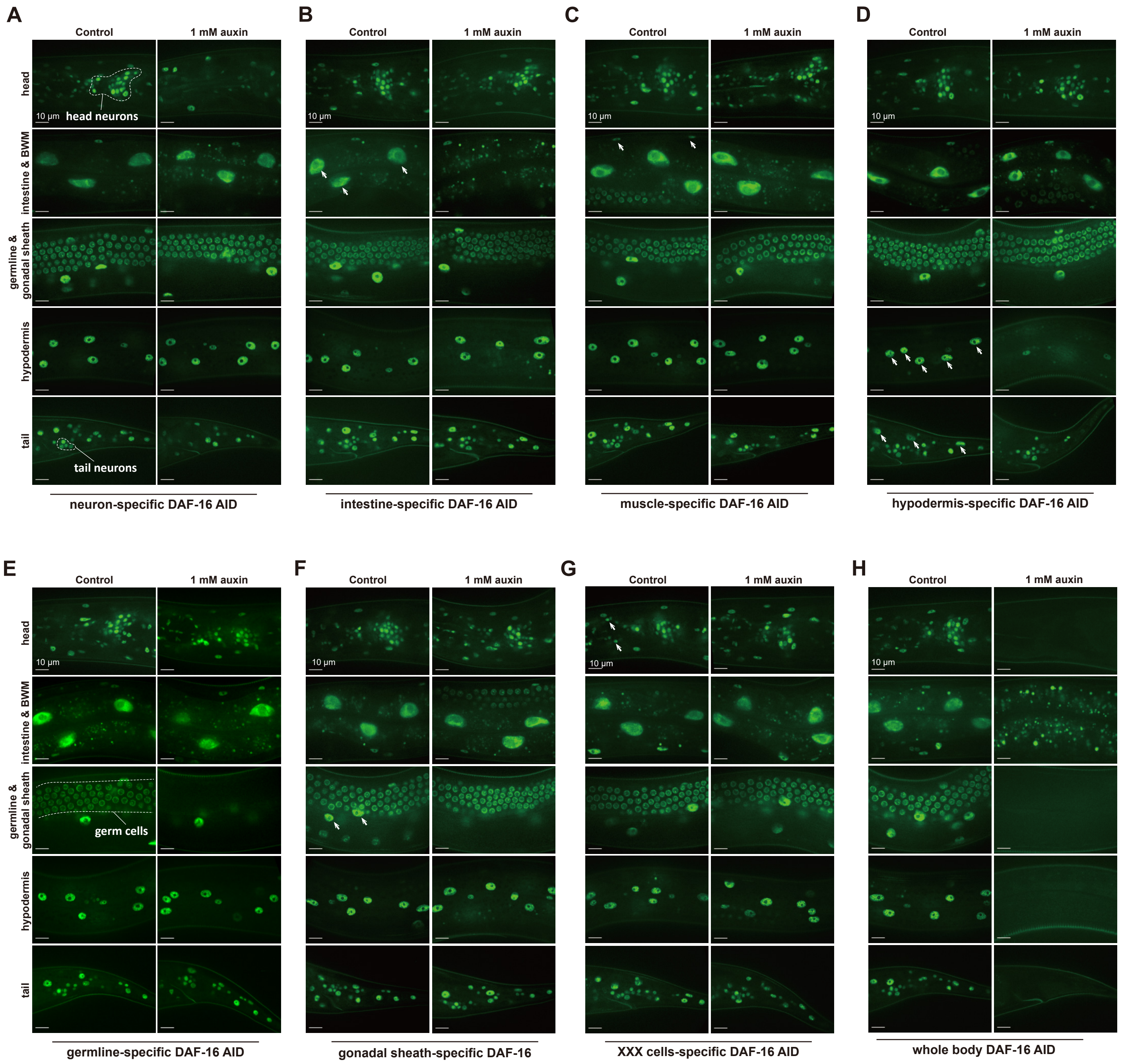
A
B
C
D
E
F
G
H
Control
1 mM auxin
head
intestine & BWM
germline & gonadal sheath
hypodermis
tail
10 μm
head neurons
tail neurons
germ cells
neuron-specific DAF-16 AID
intestine-specific DAF-16 AID
muscle-specific DAF-16 AID
hypodermis-specific DAF-16 AID
germline-specific DAF-16 AID
gonadal sheath-specific DAF-16
XXX cells-specific DAF-16 AID
whole body DAF-16 AID

### Figure S6

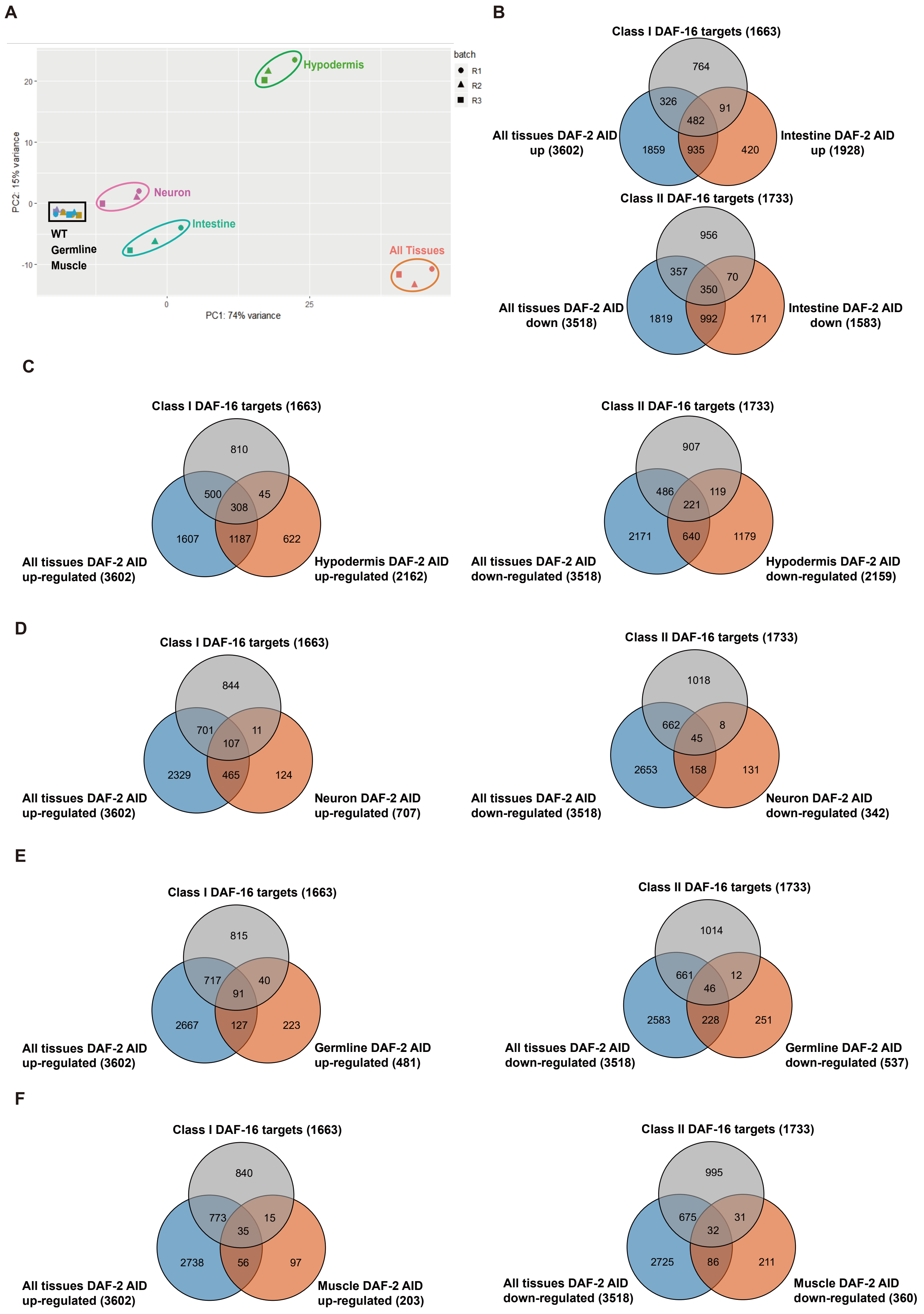
A
B
Hypodermis
Neuron
Intestine
All Tissues
WT
Germline
Muscle
batch
R1
R2
R3
PC1: 74% variance
PC2: 15% variance
Class I DAF-16 targets (1663)
764
326
91
482
1859
935
420
All tissues DAF-2 AID up (3602)
Intestine DAF-2 AID up (1928)
Class II DAF-16 targets (1733)
956
357
70
350
1819
992
171
All tissues DAF-2 AID down (3518)
Intestine DAF-2 AID down (1583)
C
Class I DAF-16 targets (1663)
810
500
45
308
1607
1187
622
All tissues DAF-2 AID up-regulated (3602)
Hypodermis DAF-2 AID up-regulated (2162)
Class II DAF-16 targets (1733)
907
486
119
221
2171
640
1179
All tissues DAF-2 AID down-regulated (3518)
Hypodermis DAF-2 AID down-regulated (2159)
D
Class I DAF-16 targets (1663)
844
701
11
107
2329
465
124
All tissues DAF-2 AID up-regulated (3602)
Neuron DAF-2 AID up-regulated (707)
Class II DAF-16 targets (1733)
1018
662
8
45
2653
158
131
All tissues DAF-2 AID down-regulated (3518)
Neuron DAF-2 AID down-regulated (342)
E
Class I DAF-16 targets (1663)
815
717
40
91
2667
127
223
All tissues DAF-2 AID up-regulated (3602)
Germline DAF-2 AID up-regulated (481)
Class II DAF-16 targets (1733)
1014
661
12
46
2583
228
251
All tissues DAF-2 AID down-regulated (3518)
Germline DAF-2 AID down-regulated (537)
F
Class I DAF-16 targets (1663)
840
773
15
35
2738
56
97
All tissues DAF-2 AID up-regulated (3602)
Muscle DAF-2 AID up-regulated (203)
Class II DAF-16 targets (1733)
995
675
31
32
2725
86
211
All tissues DAF-2 AID down-regulated (3518)
Muscle DAF-2 AID down-regulated (360)

#### Figure S7

A

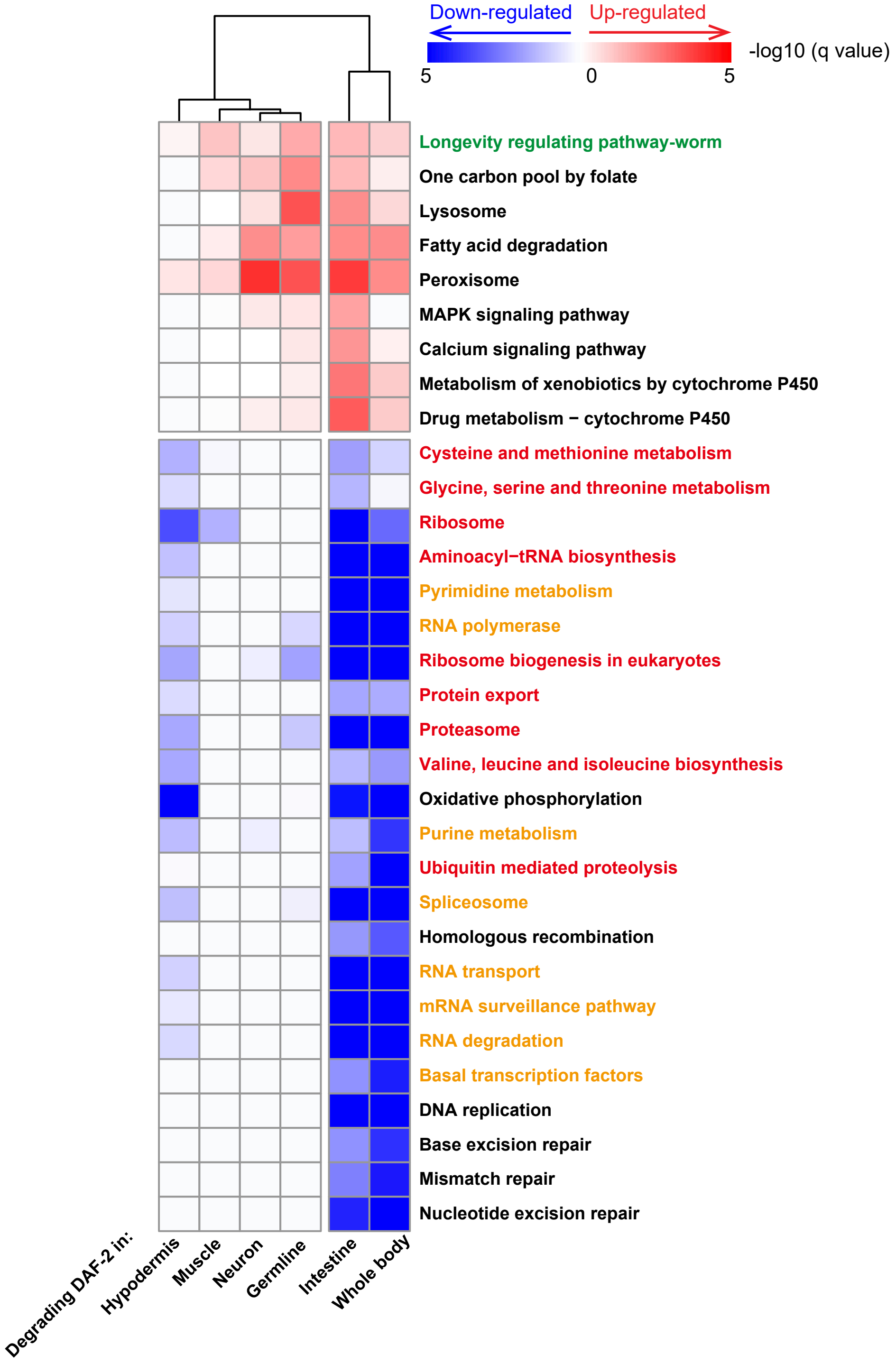

Down-regulated
Up-regulated
-log10 (q value)
5
0
5
Longevity regulating pathway-worm
One carbon pool by folate
Lysosome
Fatty acid degradation
Peroxisome
MAPK signaling pathway
Calcium signaling pathway
Metabolism of xenobiotics by cytochrome P450
Drug metabolism − cytochrome P450
Cysteine and methionine metabolism
Glycine, serine and threonine metabolism
Ribosome
Aminoacyl−tRNA biosynthesis
Pyrimidine metabolism
RNA polymerase
Ribosome biogenesis in eukaryotes
Protein export
Proteasome
Valine, leucine and isoleucine biosynthesis
Oxidative phosphorylation
Purine metabolism
Ubiquitin mediated proteolysis
Spliceosome
Homologous recombination
RNA transport
mRNA surveillance pathway
RNA degradation
Basal transcription factors
DNA replication
Base excision repair
Mismatch repair
Nucleotide excision repair
Degrading DAF-2 in:
Hypodermis
Muscle
Neuron
Germline
Intestine
Whole body


B

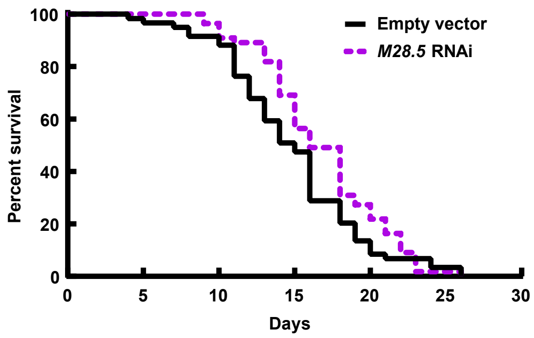

Empty vector
M28.5 RNAi
Percent survival
Days
0
20
40
60
80
100
0
5
10
15
20
25
30

### Figure S8

**A**

Wash tissue-specifc fluorecent tag labeled worms using M9

↓

Lysis buffer treatment to disrupt the cuticle

↓

Wash rapidly 5 times using M9

↓

Pronase treatment to dissociate cells

↓

20% FBS to stop cell digestion

↓

Filter to remove large debris

↓

Centrifuge to remove small debris

↓

FACS sorting fluorecent positive cells, and collect cells directly in TRiZol

**B**

| Tissue | Treatment | Replicates | Events |
|---|---|---|---|
| Intestine | Control | 4 | 8,500–20,000 |
| | Auxin | 5 | 10,000–17,500 |
| Neuron | Control | 6 | 100,000–270,000 |
| | Auxin | 6 | 160,000–280,000 |
| Hypodermis | Control | 5 | 15,000–43,500 |
| | Auxin | 6 | 19,000–35,000 |
| Muscle | Control | 4 | 7,500–12,500 |
| | Auxin | 4 | 7,500–12,300 |

**C**

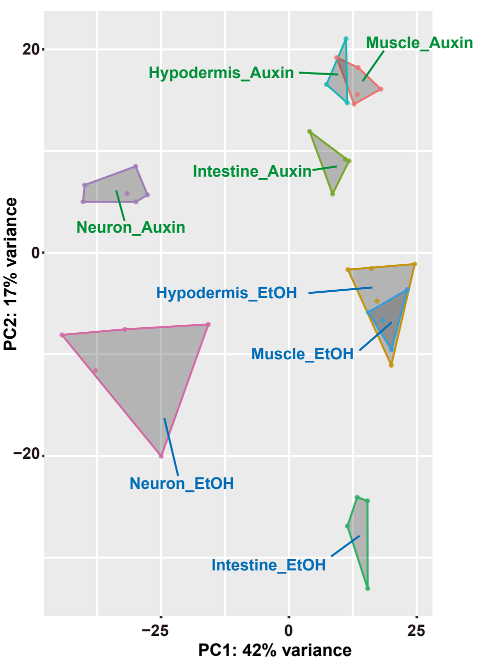


**D**

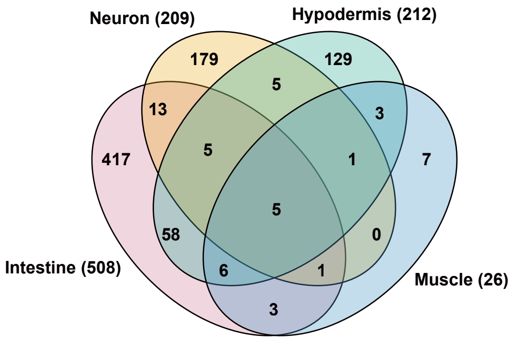


**E**

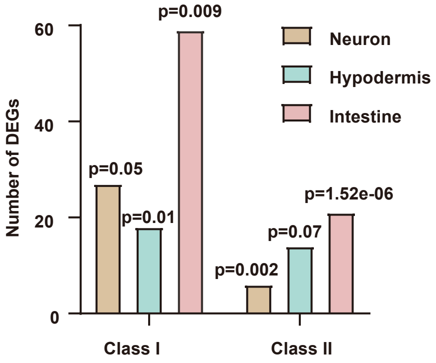


**F**

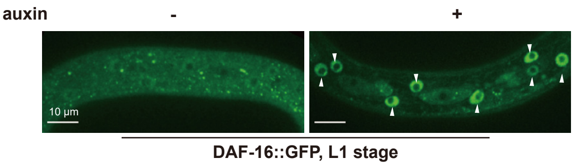


**G**

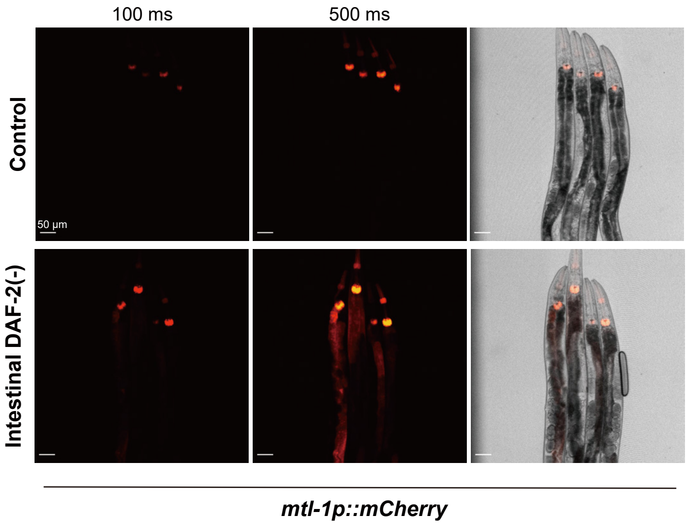


**H**

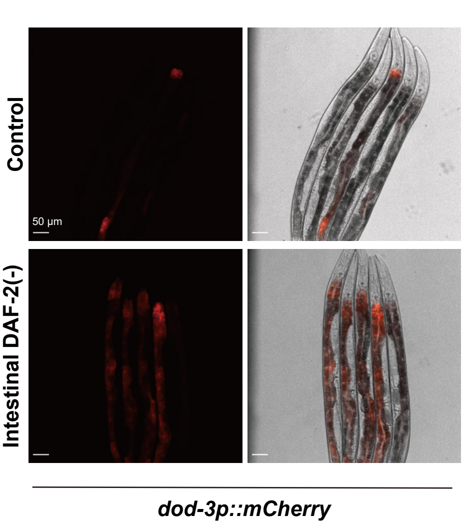

**Table S1. Statistical analyses of lifespan experiments (related to Figure 3, 4, 6 and 7)**

| Strain | Genotype | Related to | Conditions | Mean lifespan (Days) | Counted animals | Mean lifespan extension | *p*-value (Log-Rank test) |
|---|---|---|---|---|---|---|---|
| **Effects of tissue-specific DAF-2 degradation on lifespan. Lifespan assays were performed at 20 °C. Related to Figure 3.** | | | | | | | |
| MQD2356 | *hqKi373[rgef-1p::TIR1::mRuby::unc-54 3'UTR+Cbr-unc-119(+)] II; unc-119(ed3) III; daf-2(hqKi363[daf-2::degron::mNeonGreen]) III* | Figure 3A, Shown in Figure 3, Rep1 | 0.25% ethanol | 23.26 | 109 | -- | -- |
| MQD2356 | *hqKi373[rgef-1p::TIR1::mRuby::unc-54 3'UTR+Cbr-unc-119(+)] II; unc-119(ed3) III; daf-2(hqKi363[daf-2::degron::mNeonGreen]) III* | | 1 mM auxin | 27.59 | 143 | 118.62% | <0.0001 |
| MQD2375 | *unc-119(ed3) III; daf-2(hqKi363[daf-2::degron::mNeonGreen]) III; ieSi38[sun-1p::TIR1::mRuby::sun-1 3'UTR+Cbr-unc-119(+)] IV* | Figure 3B, Shown in Figure 3, Rep1 | 0.25% ethanol | 23.00 | 126 | -- | -- |
| MQD2375 | *unc-119(ed3) III; daf-2(hqKi363[daf-2::degron::mNeonGreen]) III; ieSi38[sun-1p::TIR1::mRuby::sun-1 3'UTR+Cbr-unc-119(+)] IV* | | 1 mM auxin | 24.47 | 114 | 106.41% | 0.014 |
| MQD2378 | *hqKi374[dpy-7p::TIR1::mRuby::unc-54 3'UTR+Cbr-unc-119(+)] II; unc-119(ed3) III; daf-2(hqKi363[daf-2::degron::mNeonGreen]) III* | Figure 3C, Shown in Figure 3, Rep1 | 0.25% ethanol | 22.34 | 93 | -- | -- |
| MQD2378 | *hqKi374[dpy-7p::TIR1::mRuby::unc-54 3'UTR+Cbr-unc-119(+)] II; unc-119(ed3) III; daf-2(hqKi363[daf-2::degron::mNeonGreen]) III* | | 1 mM auxin | 25.41 | 115 | 113.72% | 0.001 |

| | | | | | | | |
|---|---|---|---|---|---|---|---|
| MQD2379 | *hqKi375[myo-3p::TIR1::mRuby::unc-54 3'UTR+Cbr-unc-119(+)] II; unc-119(ed3) III; daf-2(hqKi363[daf-2::degron::mNeonGreen]) III* | Figure 3D, Shown in Figure 3, Rep1 | 0.25% ethanol | 23.32 | 82 | -- | -- |
| MQD2379 | *hqKi375[myo-3p::TIR1::mRuby::unc-54 3'UTR+Cbr-unc-119(+)] II; unc-119(ed3) III; daf-2(hqKi363[daf-2::degron::mNeonGreen]) III* | | 1 mM auxin | 24.11 | 111 | 103.39% | 0.606 |
| MQD2383 | *hqKi378[lim-7p::TIR1::mRuby::unc-54 3'UTR+Cbr-unc-119(+)] II; unc-119(ed3) III; daf-2(hqKi363[daf-2::degron::mNeonGreen]) III* | Figure 3E, Shown in Figure 3, Rep1 | 0.25% ethanol | 23.42 | 97 | -- | -- |
| MQD2383 | *hqKi378[lim-7p::TIR1::mRuby::unc-54 3'UTR+Cbr-unc-119(+)] II; unc-119(ed3) III; daf-2(hqKi363[daf-2::degron::mNeonGreen]) III* | | 1 mM auxin | 24.63 | 142 | 105.17% | 0.157 |
| MQD2402 | *unc-119(ed3) III; daf-2(hqKi363[daf-2::degron::mNeonGreen]) III; hqKi388[eak-4p::TIR-1:mRuby::unc-54 3'UTR+Cbr-unc-119(+)] IV* | Figure 3F, Shown in Figure 3, Rep1 | 0.25% ethanol | 21.21 | 97 | -- | -- |
| MQD2402 | *unc-119(ed3) III; daf-2(hqKi363[daf-2::degron::mNeonGreen]) III; hqKi388[eak-4p::TIR-1:mRuby::unc-54 3'UTR+Cbr-unc-119(+)] IV* | | 1 mM auxin | 21.01 | 106 | 99.07% | 0.529 |
| MQD2374 | *ieSi61[ges-1p::TIR1::mRuby::unc-54 3'UTR+Cbr-unc-119(+)] II; unc-119(ed3) III; daf-2(hqKi363[daf-2::degron::mNeonGreen]) III* | Figure 3G, Shown in Figure 3, Rep1 | 0.25% ethanol | 22.38 | 110 | -- | -- |
| MQD2374 | *ieSi61[ges-1p::TIR1::mRuby::unc-54 3'UTR+Cbr-unc-119(+)] II; unc-119(ed3) III; daf-2(hqKi363[daf-2::degron::mNeonGreen]) III* | | 1 mM auxin | 43.50 | 115 | 194.33% | <0.0001 |

| | | | | | | | |
|---|---|---|---|---|---|---|---|
| MQD2374 | *ieSi61[ges-1p::TIR1::mRuby::unc-54 3'UTR+Cbr-unc-119(+)] II; unc-119(ed3) III; daf-2(hqKi363[daf-2::degron::mNeonGreen]) III* | Figure 3H, Shown in Figure 3, Rep1 | 50 µg/ml FUdR, 0.25% ethanol | 21.00 | 91 | -- | -- |
| MQD2374 | *ieSi61[ges-1p::TIR1::mRuby::unc-54 3'UTR+Cbr-unc-119(+)] II; unc-119(ed3) III; daf-2(hqKi363[daf-2::degron::mNeonGreen]) III* | | 50 µg/ml FUdR, 1 mM auxin | 40.34 | 112 | 192.09% | <0.0001 |
| N2 | Wild type | | 50 µg/ml FUdR, 1 mM auxin | 20.25 | 91 | -- | -- |
| MQD855 | *daf-2(e1370ts) III* | | 50 µg/ml FUdR, 1 mM auxin | 41.71 | 113 | 205.93% | <0.0001 |
| CA1200 | *ieSi57[eft-3p::TIR-1::mRuby::unc-54 3'UTR+Cbr-unc-119(+)] II; unc-119(ed3) III* | | 50 µg/ml FUdR, 1 mM auxin | 21.19 | 95 | -- | -- |
| MQD2453 | *ieSi57[eft-3p::TIR1::mRuby::sun-1 3'UTR+Cbr-unc-119(+)] II; unc-119(ed3) III; daf-2(hqKi363[daf-2::degron::mNeonGreen]) III* | | 50 µg/ml FUdR, 1 mM auxin | 56.48 | 96 | 266.55% | <0.0001 |
| MQD2356 | *hqKi373[rgef-1p::TIR1::mRuby::unc-54 3'UTR+Cbr-unc-119(+)] II; unc-119(ed3) III; daf-2(hqKi363[daf-2::degron::mNeonGreen]) III* | Figure 3A, Rep2 | 0.25% ethanol | 22.75 | 123 | -- | -- |
| MQD2356 | *hqKi373[rgef-1p::TIR1::mRuby::unc-54 3'UTR+Cbr-unc-119(+)] II; unc-119(ed3) III; daf-2(hqKi363[daf-2::degron::mNeonGreen]) III* | | 1 mM auxin | 26.76 | 132 | 117.63% | <0.0001 |

| | | | | | | | |
|---|---|---|---|---|---|---|---|
| MQD2375 | *unc-119(ed3) III; daf-2(hqKi363[daf-2::degron::mNeonGreen]) III; ieSi38[sun-1p::TIR1::mRuby::sun-1 3'UTR+Cbr-unc-119(+)] IV* | Figure 3B, Rep2 | 0.25% ethanol | 22.03 | 130 | -- | -- |
| MQD2375 | *unc-119(ed3) III; daf-2(hqKi363[daf-2::degron::mNeonGreen]) III; ieSi38[sun-1p::TIR1::mRuby::sun-1 3'UTR+Cbr-unc-119(+)] IV* | | 1 mM auxin | 24.66 | 129 | 111.93% | <0.0001 |
| MQD2378 | *hqKi374[dpy-7p::TIR1::mRuby::unc-54 3'UTR+Cbr-unc-119(+)] II; unc-119(ed3) III; daf-2(hqKi363[daf-2::degron::mNeonGreen]) III* | Figure 3C, Rep2 | 0.25% ethanol | 22.23 | 97 | --- | -- |
| MQD2378 | *hqKi374[dpy-7p::TIR1::mRuby::unc-54 3'UTR+Cbr-unc-119(+)] II; unc-119(ed3) III; daf-2(hqKi363[daf-2::degron::mNeonGreen]) III* | | 1 mM auxin | 24.94 | 119 | 112.21% | 0.003 |
| MQD2379 | *hqKi375[myo-3p::TIR1::mRuby::unc-54 3'UTR+Cbr-unc-119(+)] II; unc-119(ed3) III; daf-2(hqKi363[daf-2::degron::mNeonGreen]) III* | Figure 3D, Rep2 | 0.25% ethanol | 22.10 | 98 | - | -- |
| MQD2379 | *hqKi375[myo-3p::TIR1::mRuby::unc-54 3'UTR+Cbr-unc-119(+)] II; unc-119(ed3) III; daf-2(hqKi363[daf-2::degron::mNeonGreen]) III* | | 1 mM auxin | 23.83 | 96 | 107.83% | 0.130 |
| MQD2383 | *hqKi378[lim-7p::TIR1::mRuby::unc-54 3'UTR+Cbr-unc-119(+)] II; unc-119(ed3) III; daf-2(hqKi363[daf-2::degron::mNeonGreen]) III* | Figure 3E, Rep2 | 0.25% ethanol | 22.71 | 103 | -- | -- |
| MQD2383 | *hqKi378[lim-7p::TIR1::mRuby::unc-54 3'UTR+Cbr-unc-119(+)] II; unc-119(ed3) III; daf-2(hqKi363[daf-2::degron::mNeonGreen]) III* | | 1 mM auxin | 23.44 | 114 | 103.21% | 0.726 |

| | | | | | | | |
|---|---|---|---|---|---|---|---|
| MQD2374 | *ieSi61[ges-1p::TIR1::mRuby::unc-54 3'UTR+Cbr-unc-119(+)] II; unc-119(ed3) III; daf-2(hqKi363[daf-2::degron::mNeonGreen]) III* | | 50 µg/ml FUdR, 0.25% ethanol | 22.63 | 105 | -- | -- |
| MQD2374 | *ieSi61[ges-1p::TIR1::mRuby::unc-54 3'UTR+Cbr-unc-119(+)] II; unc-119(ed3) III; daf-2(hqKi363[daf-2::degron::mNeonGreen]) III* | | 50 µg/ml FUdR, 1 mM auxin | 41.58 | 109 | 183.74% | <0.0001 |
| N2 | Wild type | Figure 3H, Rep2 | 50 µg/ml FUdR, 1 mM auxin | 21.16 | 93 | | -- |
| MQD855 | *daf-2(e1370ts) III* | | 50 µg/ml FUdR, 1 mM auxin | 44.73 | 91 | 211.36% | <0.0001 |
| MQD2428 | *daf-2(hqKi363[daf-2::degron::mNeonGreen]) III* | | 50 µg/ml FUdR, 1 mM auxin | 21.09 | 136 | -- | -- |
| MQD2453 | *ieSi57[eft-3p::TIR1::mRuby::sun-1 3'UTR+Cbr-unc-119(+)] II; unc-119(ed3) III; daf-2(hqKi363[daf-2::degron::mNeonGreen]) III* | | 50 µg/ml FUdR, 1 mM auxin | 55.90 | 133 | 265.03% | <0.0001 |

**Effects of tissue-specific DAF-16 degradation on the lifespan of *daf-2(e1370)* worms. Lifespan assays were performed at 20 °C. Related to Figure 4.**

| **Strain** | **Genotype** | **Related to** | **Conditions** | **Mean lifespan (Days)** | **Counted animals** | **Mean lifespan extension** | ***p*-value (Log-Rank test)** |
|---|---|---|---|---|---|---|---|

| | | | | | | | |
|---|---|---|---|---|---|---|---|
| MQD2433 | *daf-16(hqKi389[daf-16::gfp::degron]) I* | Figure 4A-4H, Shown in Figure 4, Rep1 | 1 mM auxin | 22.63 | 117 | -- | -- |
| MQD2490 | *daf-16(hqKi389[daf-16::gfp::degron]) I; daf-2(e1370ts) III* | Figure 4A-4H, Shown in Figure 4, Rep1 | 1 mM auxin | 50.95 | 114 | 225.11% | <0.0001 |
| MQD2492 | *daf-16(hqKi389[daf-16::gfp::degron]) I; hqKi373[rgef-1p::TIR1::mRuby::unc-54 3'UTR+Cbr-unc-119(+)] II; unc-119(ed3) III; daf-2(e1370ts) III* | Figure 4A, Shown in Figure 4, Rep1 | 1 mM auxin | 42.99 | 140 | 189.97% | <0.0001 |
| MQD2498 | *daf-16(hqKi389[daf-16::gfp::degron]) I; unc-119(ed3) III; daf-2(e1370ts) III; ieSi38[sun-1p::TIR1::mRuby::sun-1 3'UTR+Cbr-unc-119(+)] IV* | Figure 4B, Shown in Figure 4, Rep1 | 1 mM auxin | 47.94 | 104 | 211.83% | <0.0001 |
| MQD2493 | *daf-16(hqKi389[daf-16::gfp::degron]) I; unc-119(ed3) III; hqKi374[dpy-7p::TIR1::mRuby::unc-54 3'UTR+Cbr-unc-119(+)] II; daf-2(e1370ts) III* | Figure 4C, Shown in Figure 4, Rep1 | 1 mM auxin | 43.02 | 124 | 190.10% | <0.0001 |
| MQD2499 | *daf-16(hqKi389[daf-16::gfp::degron]) I; hqKi375[myo-3p::TIR1::mRuby::unc-54 3'UTR+Cbr-unc-119(+)] II; unc-119(ed3) III; daf-2(e1370ts) III* | Figure 4D, Shown in Figure 4, Rep1 | 1 mM auxin | 48.54 | 120 | 214.48% | <0.0001 |
| MQD2500 | *daf-16(hqKi389[daf-16::gfp::degron]) I; hqKi378[lim-7p::TIR1::mRuby::unc-54 3'UTR+Cbr-unc-119(+)] II; unc-119(ed3) III; daf-2(e1370ts) III* | Figure 4E, Shown in Figure 4, Rep1 | 1 mM auxin | 49.75 | 120 | 219.82% | <0.0001 |

| | | | | | | | |
|---|---|---|---|---|---|---|---|
| MQD2495 | *daf-16(hqKi389[Pdaf-16::daf-16::gfp::degron]) I; unc-119(ed3) III; daf-2(e1370ts) III; hqKi388[Peak-4::TIR-1:mRuby::unc-54 3'UTR+Cbr-unc-119(+)] IV* | Figure 4F, Shown in Figure 4, Rep1 | 1 mM auxin | 48.92 | 101 | 216.16% | <0.0001 |
| MQD2494 | *daf-16(hqKi389[daf-16::gfp::degron]) I; ieSi61[ges-1p::TIR1::mRuby::unc-54 3'UTR+Cbr-unc-119(+)] II; unc-119(ed3) III; daf-2(e1370ts) III* | Figure 4G, Shown in Figure 4, Rep1 | 1 mM auxin | 30.50 | 127 | 134.78% | <0.0001 |
| MQD2491 | *daf-16(hqKi389[daf-16::gfp::degron]) I; ieSi57[eft-3p::TIR1::mRuby::unc-54 3'UTR+Cbr-unc-119(+)] II; unc-119(ed3) III; daf-2(e1370ts) III* | Figure 4H, Shown in Figure 4, Rep1 | 1 mM auxin | 21.50 | 117 | 95.02% | 0.043 |
| **Lifespan assays were performed at 25 °C.** | | | | | | | |
| MQD2433 | *daf-16(hqKi389[daf-16::gfp::degron]) I* | Figure 4, Rep2 | 1 mM auxin | 15.74 | 113 | -- | -- |
| MQD2490 | *daf-16(hqKi389[daf-16::gfp::degron]) I; daf-2(e1370ts) III* | Figure 4, Rep2 | 1 mM auxin | 35.16 | 141 | 223.47% | <0.0001 |
| MQD2492 | *daf-16(hqKi389[daf-16::gfp::degron]) I; hqKi373[rgef-1p::TIR1::mRuby::unc-54 3'UTR+Cbr-unc-119(+)] II; unc-119(ed3) III; daf-2(e1370ts) III* | Figure 4A, Rep2 | 1 mM auxin | 30.15 | 130 | 191.64% | <0.0001 |
| MQD2498 | *daf-16(hqKi389[daf-16::gfp::degron]) I; unc-119(ed3) III; daf-2(e1370ts) III; ieSi38[sun-1p::TIR1::mRuby::sun-1 3'UTR+Cbr-unc-119(+)] IV* | Figure 4B, Rep2 | 1 mM auxin | 31.76 | 123 | 201.82% | <0.0001 |

| | | | | | | | |
|---|---|---|---|---|---|---|---|
| MQD2493 | *daf-16(hqKi389[daf-16::gfp::degron]) I; unc-119(ed3) III; hqKi374[dpy-7p::TIR1::mRuby::unc-54 3'UTR+Cbr-unc-119(+)] II; daf-2(e1370ts) III* | Figure 4C, Rep2 | 1 mM auxin | 30.43 | 120 | 193.39% | <0.0001 |
| MQD2499 | *daf-16(hqKi389[daf-16::gfp::degron]) I; hqKi375[myo-3p::TIR1::mRuby::unc-54 3'UTR+Cbr-unc-119(+)] II; unc-119(ed3) III; daf-2(e1370ts) III* | Figure 4D, Rep2 | 1 mM auxin | 33.61 | 134 | 213.60% | <0.0001 |
| MQD2500 | *daf-16(hqKi389[daf-16::gfp::degron]) I; hqKi378[lim-7p::TIR1::mRuby::unc-54 3'UTR+Cbr-unc-119(+)] II; unc-119(ed3) III; daf-2(e1370ts) III* | Figure 4E, Rep2 | 1 mM auxin | 34.26 | 125 | 217.71% | <0.0001 |
| MQD2494 | *daf-16(hqKi389[daf-16::gfp::degron]) I; ieSi61[ges-1p::TIR1::mRuby::unc-54 3'UTR+Cbr-unc-119(+)] II; unc-119(ed3) III; daf-2(e1370ts) III* | Figure 4G, Rep2 | 1 mM auxin | 18.96 | 127 | 120.50% | <0.0001 |
| MQD2491 | *daf-16(hqKi389[daf-16::gfp::degron]) I; ieSi57[eft-3p::TIR1::mRuby::unc-54 3'UTR+Cbr-unc-119(+)] II; unc-119(ed3) III; daf-2(e1370ts) III* | Figure 4H, Rep2 | 1 mM auxin | 14.45 | 116 | 91.82% | 0.659 |

**Effects of intestine-specific DAF-16 degradation on the lifespan of intestinal DAF-2 AID worms. Lifespan assays were performed at 20 °C. Related to Figure 4J**

| Strain | Genotype | Related to | Conditions | Mean lifespan (Days) | Counted animals | Mean lifespan extension | *p*-value (Log-Rank test) |
|---|---|---|---|---|---|---|---|

| | | | | | | | |
|---|---|---|---|---|---|---|---|
| MQD2374 | *ieSi61[ges-1p::TIR1::mRuby::unc-54 3'UTR+Cbr-unc-119(+)] II; unc-119(ed3) III; daf-2(hqKi363[daf-2::degron::mNeonGreen]) III* | Figure 4J (Shown in the Figure 4), Rep1 | 0.25% ethanol | 23.48 | 124 | -- | -- |
| MQD2374 | *ieSi61[ges-1p::TIR1::mRuby::unc-54 3'UTR+Cbr-unc-119(+)] II; unc-119(ed3) III; daf-2(hqKi363[daf-2::degron::mNeonGreen]) III* | | 1 mM auxin | 46.03 | 96 | 196.01% | <0.0001 |
| MQD2477 | *daf-16(hqKi389[Pdaf-16::daf-16::gfp::degron]) I; ieSi61[ges-1p::TIR1::mRuby::unc-54 3'UTR+ Cbr-unc-119(+)] II; unc-119(ed3) III; daf-2(hqKi363[daf-2::degron::mNeonGreen]) III* | | 0.25% ethanol | 21.98 | 104 | -- | -- |
| MQD2477 | *daf-16(hqKi389[Pdaf-16::daf-16::gfp::degron]) I; ieSi61[ges-1p::TIR1::mRuby::unc-54 3'UTR+ Cbr-unc-119(+)] II; unc-119(ed3) III; daf-2(hqKi363[daf-2::degron::mNeonGreen]) III* | | 1 mM auxin | 27.20 | 120 | 123.74% | <0.0001 |
| MQDD2374 | *ieSi61[ges-1p::TIR1::mRuby::unc-54 3'UTR+Cbr-unc-119(+)] II; unc-119(ed3) III; daf-2(hqKi363[daf-2::degron::mNeonGreen]) III* | Figure 4J | 0.25% ethanol | 22.40 | 101 | -- | -- |
| MQD2374 | *ieSi61[ges-1p::TIR1::mRuby::unc-54 3'UTR+Cbr-unc-119(+)] II; unc-119(ed3) III; daf-2(hqKi363[daf-2::degron::mNeonGreen]) III* | | 1 mM auxin | 42.39 | 104 | 189.25% | <0.0001 |
| MQD2477 | *daf-16(hqKi389[Pdaf-16::daf-16::gfp::degron]) I; ieSi61[ges-1p::TIR1::mRuby::unc-54 3'UTR+ Cbr-unc-119(+)] II; unc-119(ed3) III; daf-2(hqKi363[daf-2::degron::mNeonGreen]) III* | | 0.25% ethanol | 21.42 | 125 | -- | -- |

| | | | | | | | |
|---|---|---|---|---|---|---|---|
| MQD2477 | *daf-16(hqKi389[Pdaf-16::daf-16::gfp::degron]) I; ieSi61[ges-1p::TIR1::mRuby::unc-54 3'UTR+ Cbr-unc-119(+)] II; unc-119(ed3) III; daf-2(hqKi363[daf-2::degron::mNeonGreen]) III* | | 1 mM auxin | 25.17 | 116 | 117.49% | <0.0001 |

**Effects of *fib-1* RNAi and *M28.5* RNAi on the lifespan of *daf-2(e1370)* worms. Lifespan assays were performed at 25 °C. Related to Figure 6D**

| Strain | Genotype | Related to | Conditions | Mean lifespan (Days) | Counted animals | Mean lifespan extension | *p*-value (Log-Rank test) |
|---|---|---|---|---|---|---|---|
| MQD855 | *daf-2(e1370ts) III* | | empty vector | 30.66 | 61 | -- | -- |
| MQD855 | *daf-2(e1370ts) III* | Figure 6D | *fib-1* RNAi | 35.38 | 97 | 115.39% | 0.005 |
| MQD855 | *daf-2(e1370ts) III* | | *M28.5* RNAi | 35.96 | 93 | 117.29% | 0.001 |

**Effects of *M28.5* RNAi on the lifespan of N2 worms. Lifespan assays were performed at 25 °C. Related to Figure S7B**

| Strain | Genotype | Related to | Conditions | Mean lifespan (Days) | Counted animals | Mean lifespan extension | *p*-value (Log-Rank test) |
|---|---|---|---|---|---|---|---|
| N2 | Wild type | | empty vector | 14.62 | 61 | -- | -- |
| N2 | Wild type | Figure S7B | *M28.5* RNAi | 16.52 | 52 | 113.00% | 0.092 |

**Effects of simultaneous degradation of intestinal DAF-2 and non-intestinal DAF-16. Lifespan assays were performed at 20 °C. Related to Figure 7F.**

| Strain | Genotype | Related to | Conditions | Mean lifespan (Days) | Counted animals | Mean lifespan extension | *p*-value (Log-Rank test) |
|---|---|---|---|---|---|---|---|
| MQD1508 | *daf-2(hqKi14[daf-2::gfp]) III* | Figure 7F, top panel, shown in the Figure | 20 °C | 21.29 | 118 | -- | -- |
| OD2768 | *ItSi910[elt-2p::vhhGFP4::zif-1::operon-linker::mCherry::his-11::tbb-2 3'UTR+Cbr-unc-119(+)] II; unc-119(ed3) III* | | | 20.54 | 114 | 96.48% | 0.271 |
| MQD2778 | *ItSi910[elt-2p::vhhGFP4::zif-1::operon-linker::mCherry::his-11::tbb-2 3'UTR+Cbr-unc-119(+)] II; unc-119(ed3) III; daf-2(hqKi14[daf-2::gfp]) III* | | | 31.82 | 114 | 149.46% | <0.0001 |
| MQD2783 | *daf-16(hqKi490[daf-16::tagBFP::degron]) I; ItSi910[elt-2p::vhhGFP4::zif-1::operon-linker::mCherry::his-11::tbb-2 3'UTR+Cbr-unc-119(+)] II; unc-119(ed3) III; daf-2(hqKi14[daf-2::gfp]) III; hqKi403[rgef-1p::TIR-1::mRuby::unc-54 3'UTR+Cbr-unc-119(+)]) IV* | Figure 7F, middle panel, shown in the Figure, Rep1 | 0.25% ethanol | 30.41 | 128 | -- | -- |
| MQD2783 | *daf-16(hqKi490[daf-16::tagBFP::degron]) I; ItSi910[elt-2p::vhhGFP4::zif-1::operon-linker::mCherry::his-11::tbb-2 3'UTR+Cbr-unc-119(+)] II; unc-119(ed3) III; daf-2(hqKi14[daf-2::gfp]) III; hqKi403[rgef-1p::TIR-1::mRuby::unc-54 3'UTR+Cbr-unc-119(+)]) IV* | | 1 mM auxin | 30.66 | 129 | 100.82% | 0.337 |

| | | | | | | | |
|---|---|---|---|---|---|---|---|
| MQD2781 | *daf-16(hqKi488[daf-16::tagBFP::degron]) I; ItSi910[elt-2p::vhhGFP4::zif-1::operon-linker::mCherry::his-11::tbb-2 3'UTR+Cbr-unc-119(+)]) II; unc-119(ed3) III; daf-2(hqKi14[daf-2::gfp]) III; hqKi407[dpy-7p::TIR-1::mRuby::unc-54 3'UTR+Cbr-unc-119(+)] IV* | Figure 7F, bottom panel,shown in the Figure, Rep1 | 0.25% ethanol | 31.56 | 119 | -- | -- |
| MQD2781 | *daf-16(hqKi488[daf-16::tagBFP::degron]) I; ItSi910[elt-2p::vhhGFP4::zif-1::operon-linker::mCherry::his-11::tbb-2 3'UTR+Cbr-unc-119(+)]) II; unc-119(ed3) III; daf-2(hqKi14[daf-2::gfp]) III; hqKi407[dpy-7p::TIR-1::mRuby::unc-54 3'UTR+Cbr-unc-119(+)] IV* | | 1 mM auxin | 28.17 | 113 | 89.26% | 0.003 |
| MQD2783 | *daf-16(hqKi490[daf-16::tagBFP::degron]) I; ItSi910[elt-2p::vhhGFP4::zif-1::operon-linker::mCherry::his-11::tbb-2 3'UTR+Cbr-unc-119(+)] II; unc-119(ed3) III; daf-2(hqKi14[daf-2::gfp]) III; hqKi403[rgef-1p::TIR-1::mRuby::unc-54 3'UTR+Cbr-unc-119(+)]) IV* | Figure 7F, middle panel, Rep2 | 0.25% ethanol | 30.96 | 108 | -- | -- |
| MQD2783 | *daf-16(hqKi490[daf-16::tagBFP::degron]) I; ItSi910[elt-2p::vhhGFP4::zif-1::operon-linker::mCherry::his-11::tbb-2 3'UTR+Cbr-unc-119(+)] II; unc-119(ed3) III; daf-2(hqKi14[daf-2::gfp]) III; hqKi403[rgef-1p::TIR-1::mRuby::unc-54 3'UTR+Cbr-unc-119(+)]) IV* | | 1 mM auxin | 30.18 | 114 | 97.48% | 0.365 |

| | | | | | | | |
|---|---|---|---|---|---|---|---|
| MQD2781 | *daf-16(hqKi488[daf-16::tagBFP::degron]) I; ItSi910[elt-2p::vhhGFP4::zif-1::operon-linker::mCherry::his-11::tbb-2 3'UTR+Cbr-unc-119(+)]) II; unc-119(ed3) III; daf-2(hqKi14[daf-2::gfp]) III; hqKi407[dpy-7p::TIR-1::mRuby::unc-54 3'UTR+Cbr-unc-119(+)] IV* | Figure 7F, bottom panel, Rep2 | 0.25% ethanol | 30.88 | 102 | -- | -- |
| MQD2781 | *daf-16(hqKi488[daf-16::tagBFP::degron]) I; ItSi910[elt-2p::vhhGFP4::zif-1::operon-linker::mCherry::his-11::tbb-2 3'UTR+Cbr-unc-119(+)]) II; unc-119(ed3) III; daf-2(hqKi14[daf-2::gfp]) III; hqKi407[dpy-7p::TIR-1::mRuby::unc-54 3'UTR+Cbr-unc-119(+)] IV* | | 1 mM auxin | 28.44 | 107 | 92.10% | 0.027 |

**DAF-16::GFP or DAF-2::mNeonGreen knocking in does not affect WT lifespan. Lifespan assays were performed at 20 °C. Related to Figure S1A.**

| Strain | Genotype | Related to | Conditions | Mean lifespan (Days) | Counted animals | Mean lifespan extension | *p*-value (Log-Rank test) |
|---|---|---|---|---|---|---|---|
| N2 | Wild type | Figure S1A | - | 22.47 | 76 | -- | -- |
| MQD1543 | *daf-16(hqKi23[daf-16::gfp])* | | | 22.97 | 69 | 102.23% | 0.883 |
| MQD1661 | *daf-2(hq61Ki[daf-2::mNeongreen])* | | | 22.12 | 114 | 98.44% | 0.899 |

**Table S2 Strains used in this study**

| Strain name | Genotype | Source |
|---|---|---|
| N2 | | CGC |
| MQD855 | *daf-2(e1370ts) III* | This study |
| MQD1543 | *daf-16(hqKi23[daf-16::gfp])* | This study |
| MQD1661 | *daf-2(hq61Ki[daf-2::mNeongreen])* | This study |
| MQD1779 | *daf-2(hqKi63[daf-2::icr::NLS::gfp::mNeonGreen::NLS])* | This study |
| MQD2256 | *daf-2(hqKi63[daf-2::icr::NLS::gfp::mNeonGreen::NLS]) III; hqEx536[lim-7p::mCherry::NLS+rol-6(su1006)]* | This study |
| MQD2428 | *daf-2(hqKi363[daf-2::degron::mNeonGreen]) III* | This study |
| MQD2374 | *ieSi61[ges-1p::TIR1::mRuby::unc-54 3'UTR+Cbr-unc-119(+)] II; unc-119(ed3) III; daf-2(hqKi363[daf-2::degron::mNeonGreen]) III* | This study |
| MQD2356 | *hqKi373[rgef-1p::TIR1::mRuby::unc-54 3'UTR+Cbr-unc-119(+)] II; unc-119(ed3) III; daf-2(hqKi363[daf-2::degron::mNeonGreen]) III* | This study |
| MQD2375 | *unc-119(ed3) III; daf-2(hqKi363[daf-2::degron::mNeonGreen]) III; ieSi38[sun-1p::TIR1::mRuby::sun-1 3'UTR+Cbr-unc-119(+)] IV* | This study |
| MQD2378 | *hqKi374[dpy-7p::TIR1::mRuby::unc-54 3'UTR+Cbr-unc-119(+)] II; unc-119(ed3) III; daf-2(hqKi363[daf-2::degron::mNeonGreen]) III* | This study |
| MQD2379 | *hqKi375[myo-3p::TIR1::mRuby::unc-54 3'UTR+Cbr-unc-119(+)] II; unc-119(ed3) III; daf-2(hqKi363[daf-2::degron::mNeonGreen]) III* | This study |
| MQD2383 | *hqKi378[lim-7p::TIR1::mRuby::unc-54 3'UTR+Cbr-unc-119(+)] II; unc-119(ed3) III; daf-2(hqKi363[daf-2::degron::mNeonGreen]) III* | This study |
| MQD2453 | *ieSi57[eft-3p::TIR1::mRuby::unc-54 3'UTR+Cbr-unc-119(+)] II; unc-119(ed3) III; daf-2(hqKi363[daf-2::degron::mNeonGreen]) III* | This study |
| MQD2402 | *unc-119(ed3) III; daf-2(hqKi363[daf-2::degron::mNeonGreen]) III; hqKi388[eak-4p::TIR-1:mRuby::unc-54 3'UTR+Cbr-unc-119(+)] IV* | This study |

| Strain name | Genotype | Source |
|---|---|---|
| CA1200 | *ieSi57[eft-3p::TIR-1::mRuby::unc-54 3'UTR+Cbr-unc-119(+)] II; unc-119(ed3) III* | CGC |
| CA1199 | *unc-119(ed3) III; ieSi38[sun-1p::TIR1::mRuby::sun-1 3'UTR+ Cbr-unc-119(+)] IV* | CGC |
| MQD2490 | *daf-16(hqKi389[daf-16::gfp::degron]) I; daf-2(e1370ts) III* | This study |
| MQD2494 | *daf-16(hqKi389[daf-16::gfp::degron]) I; ieSi61[ges-1p::TIR1::mRuby::unc-54 3'UTR+Cbr-unc-119(+)] II; unc-119(ed3) III; daf-2(e1370ts) III* | This study |
| MQD2492 | *daf-16(hqKi389[daf-16::gfp::degron]) I; hqKi373[rgef-1p::TIR1::mRuby::unc-54 3'UTR+Cbr-unc-119(+)] II; unc-119(ed3) III; daf-2(e1370ts) III* | This study |
| MQD2498 | *daf-16(hqKi389[daf-16::gfp::degron]) I; unc-119(ed3) III; daf-2(e1370ts) III; ieSi38[sun-1p::TIR1::mRuby::sun-1 3'UTR+Cbr-unc-119(+)] IV* | This study |
| MQD2493 | *daf-16(hqKi389[daf-16::gfp::degron]) I; unc-119(ed3) III; hqKi374[dpy-7p::TIR1::mRuby::unc-54 3'UTR+Cbr-unc-119(+)] II; daf-2(e1370ts) III* | This study |
| MQD2499 | *daf-16(hqKi389[daf-16::gfp::degron]) I; hqKi375[myo-3p::TIR1::mRuby::unc-54 3'UTR+Cbr-unc-119(+)] II; unc-119(ed3) III; daf-2(e1370ts) III* | This study |
| MQD2500 | *daf-16(hqKi389[daf-16::gfp::degron]) I; hqKi378[lim-7p::TIR1::mRuby::unc-54 3'UTR+Cbr-unc-119(+)] II; unc-119(ed3) III; daf-2(e1370ts) III* | This study |
| MQD2491 | *daf-16(hqKi389[daf-16::gfp::degron]) I; ieSi57[eft-3p::TIR1::mRuby::unc-54 3'UTR+Cbr-unc-119(+)] II; unc-119(ed3) III; daf-2(e1370ts) III* | This study |
| MQD2433 | *daf-16(hqKi389[daf-16::gfp::degron]) I* | This study |
| MQD2495 | *daf-16(hqKi389[Pdaf-16::daf-16::gfp::degron]) I; unc-119(ed3) III; daf-2(e1370ts) III; hqKi388[Peak-4::TIR-1:mRuby::unc-54 3'UTR+Cbr-unc-119(+)] IV* | This study |
| MQD2477 | *daf-16(hqKi389[Pdaf-16::daf-16::gfp::degron]) I; ieSi61[ges-1p::TIR1::mRuby::unc-54 3'UTR+ Cbr-unc-119(+)] II; unc-119(ed3) III; daf-2(hqKi363[daf-2::degron::mNeonGreen]) III* | This study |

| Strain name | Genotype | Source |
|---|---|---|
| MQD2476 | *daf-16(hqKi23[daf-16::gfp::6his]) I; ieSi61[ges-1p::TIR1::mRuby::unc-54 3'UTR+ Cbr-unc-119(+)] II; unc-119(ed3) III; daf-2(hqKi363[daf-2::degron::mNeonGreen]) III* | This study |
| MQD2714 | *ieSi61[ges-1p::TIR1::mRuby::unc-54 3'UTR+ Cbr-unc-119(+)] II; unc-119(ed3) III; daf-2(hqKi363[daf-2::degron::mNeonGreen]) III; hqEx601[rgef-1p::NLS::gfp::mNeonGreen::NLS] .* | This study |
| MQD2726 | *ieSi61[ges-1p::TIR1::mRuby::unc-54 3'UTR+ Cbr-unc-119(+)] II; unc-119(ed3) III; daf-2(hqKi363[daf-2::degron::mNeonGreen]) III; hqEx603[ges-1p::NLS::gfp::mNeonGreen::NLS]* | This study |
| MQD2927 | *ieSi61[ges-1p::TIR1::mRuby::unc-54 3'UTR+ Cbr-unc-119(+)] II; unc-119(ed3) III; daf-2(hqKi363[daf-2::degron::mNeonGreen]) III; hqEx611[dpy-7p::NLS::gfp::NLS]* | This study |
| MQD2741 | *ieSi61[ges-1p::TIR1::mRuby::unc-54 3'UTR+ Cbr-unc-119(+)] II; unc-119(ed3) III; daf-2(hqKi363[daf-2::degron::mNeonGreen]) III; hqEx626[myo-3p::NLS::mNeonGreen::GFP::NLS]* | This study |
| MQD2755 | *ieSi61[ges-1p::TIR1::mRuby::unc-54 3'UTR+ Cbr-unc-119(+)] II; unc-119(ed3) III; daf-2(hqKi363[daf-2::degron::mNeonGreen]) III; hqEx629[mtl-1p::mCherry]* | This study |
| MQD2761 | *ieSi61[ges-1p::TIR1::mRuby::unc-54 3'UTR+ Cbr-unc-119(+)] II; unc-119(ed3) III; daf-2(hqKi363[daf-2::degron::mNeonGreen]) III; hqEx633[nas-28p::mCherry+rol-6(su1006)]* | This study |
| MQD2762 | *ieSi61[ges-1p::TIR1::mRuby::unc-54 3'UTR+ Cbr-unc-119(+)] II; unc-119(ed3) III; daf-2(hqKi363[daf-2::degron::mNeonGreen]) III; hqEx634[dod-3p::mCherry+rol-6(su1006)]* | This study |
| MQD2769 | *ieSi61[ges-1p::TIR1::mRuby::unc-54 3'UTR+ Cbr-unc-119(+)] II; daf-2(hqKi363[daf-2::degron::mNeonGreen]) III; hqEx641[gst-30p:mCherry+rol-6(su1006)]* | This study |
| MQD2771 | *ieSi61[ges-1p::TIR1::mRuby::unc-54 3'UTR+Cbr-unc-119(+)] II; unc-119(ed3) III; daf-2(hqKi363[daf-2::degron::mNeonGreen]) III; hqEx643[col-185p::mCherry+rol-6(su1006)]* | This study |

| Strain name | Genotype | Source |
|---|---|---|
| MQD1508 | *daf-2(hqKi14[daf-2::gfp]) III* | This study |
| OD2768 | *ItSi910[elt-2p::vhhGFP4::zif-1::operon-linker::mCherry::his-11::tbb-2 3'UTR+Cbr-unc-119(+)] II; unc-119(ed3) III* | CGC |
| MQD2778 | *ItSi910[elt-2p::vhhGFP4::zif-1::operon-linker::mCherry::his-11::tbb-2 3'UTR+Cbr-unc-119(+)] II; unc-119(ed3) III; daf-2(hqKi14[daf-2::gfp]) III* | This study |
| MQD2781 | *daf-16(hqKi488[daf-16::tagBFP::degron]) I; ItSi910[elt-2p::vhhGFP4::zif-1::operon-linker::mCherry::his-11::tbb-2 3'UTR+Cbr-unc-119(+)]) II; unc-119(ed3) III; daf-2(hqKi14[daf-2::gfp]) III; hqKi407[dpy-7p::TIR-1::mRuby::unc-54 3'UTR+Cbr-unc-119(+)] IV* | This study |
| MQD2783 | *daf-16(hqKi490[daf-16::tagBFP::degron]) I; ItSi910[elt-2p::vhhGFP4::zif-1::operon-linker::mCherry::his-11::tbb-2 3'UTR+Cbr-unc-119(+)] II; unc-119(ed3) III; daf-2(hqKi14[daf-2::gfp]) III; hqKi403[rgef-1p::TIR-1::mRuby::unc-54 3'UTR+Cbr-unc-119(+)]) IV* | This study |

**Table S3 Oligonucleotide sequences used in this study**

| Name | Sequence (5'-3') |
|---|---|
| **sgRNA sequences for constructing knock-in strains** | |
| sgRNAs for knocking-in the *gfp, gfp::degron*, or *tagBFP::degron* at the C-terminus of *daf-16* | |
| daf-16-sg1 | TCTCTTTCGAACAACACCAG |
| daf-16-sg2 | TCTCTTCATTTTGTTTCCCC |
| sgRNAs for knocking-in the *mNeonGreen, icr::NuGFP, degron::mNeonGreen, or gfp* at the C-terminus of *daf-2* | |
| daf-2-sg1 | TTCAACGGACGTTCGGCTTT |
| daf-2-sg2 | TTTTGGGGGTTTCAGACAAG |
| daf-2-sg3 | TGACATTTTCAACGGACGTT |
| sgRNAs for replacing the *eft-3* promoter by *rgef-1*, *dpy-7*, *myo-3*, or *lim-7* promoter on chromosome II of CA1200 strain | |
| rgef-1-sg1 | CGTGGATCCAGATATCCTGC |
| rgef-1-sg2 | CGTCACCATGGGAAGCTTCG |
| rgef-1-sg3 | TACTTCTTCTGGAAACGACA |
| sgRNAs for replacing the *sun-1* promoter by *eak-4* promoter on chromosome IV of CA1199 strain | |
| eak-4-sg1 | GGGTTAATACGACTCACTAG |
| eak-4-sg2 | Same as rgef-1-sg3 |
| **qRT-PCR primers related to Figure 6C** | |
| act-1-F | TGCCGCTCTTGTTGTAGACAATGG |
| act-1-R | TGACGTGGTCTTCCGACAATGGAT |
| pmp-3-F | TGGCCGGATGATGGTGTCGC |
| pmp-3-R | ACGAACAATGCCAAAGGCCAGC |
| 18S rRNA-F | ATGGCCGTTCTTAGTTGGTGGAGT |
| 18S rRNA-R | TATCGCTCAATCTCGTGCGGCTAA |
| 28S rRNA-F | TTGGCCAGTTGGTTGATGCTTGTC |
| 28S rRNA-R | AGCACAATCACTAGTCCGCCATCA |
| 5S rRNA-F | CGACCATATCACGTTGAATGCA |
| 5S rRNA-R | GGCCGTCTCCGATCCAAGTA |
| ITS-F | CGAAATGTCAACGTTCCAGTTG |
| ITS-R | CAACGTCTACTGCAGAACAGCAA |
| ITS2-F | TTCGCGTCTCGGCATACTG |
| ITS2-R | TCTTGATGATCAGCCGAGTCAA |
